# Multilocus typing of *Lachancea thermotolerans* for wine fermentation monitoring

**DOI:** 10.1101/2022.12.12.518888

**Authors:** Javier Vicente, Eva Navascués, Santiago Benito, Domingo Marquina, Antonio Santos

## Abstract

Climate change is causing a lack of acidity during winemaking and oenologists use several solutions to cope with such a problem. *Lachancea thermotolerans*, which has the potential to tolerate the harsh physicochemical conditions of wine, has emerged as a promising alternative for pH management during winemaking and, currently, it is the most valuable yeast used for acidity control in wine. In this work an amenable method for *L. thermotolerans* genotyping based on a multiplexed microsatellite amplification in 6 different loci was developed. This specific and sensitive method was used to distinguish between 103 collection strains obtained from different geographical and isolation sources, and then challenged against a 429 *L. thermotolerans* isolates from several wineries and harvests. The procedure was also tested for fermentation monitoring and strain implantation. The procedure was conceived to simplify the methodology available for *L. thermotolerans* genotyping, making it easy for applying in wine-related laboratories. This method can be applied to distinguish between *L. thermotolerans* strains in selection programs and to follow implantation of inoculated strains during winemaking with optimal results.

## 1. INTRODUCTION

The new climatic conditions in the different wine-growing areas due to global climate change are altering the composition of the wine grape, triggering consequences in the process of wine production (Santillán *et al*. 2019). Rising temperatures brings changes in vine phenology causing modifications in the chemical composition and microbiological quality of the grapes that arrives at the winery. In this regard, the main grape changes are related to higher sugars concentration (implying higher probable alcoholic degrees) and a lower acidity. Microbial populations developed during the winemaking process and sensory quality of the wine are affected by these changes (De Orduña 2010; Volschenk *et al*. 2006).

There are several tools for acidity control in wines based on acid or base addition or the use of different microorganisms (Vicente *et al*., 2022). There are some mechanisms to control the acidity of must and wine authorized by the International Wine Organization (OIV, 2021; Volschenk *et al*. 2006). Nevertheless, numerous studies show the interest of using selected yeast strains for wine pH regulation (Pacheco *et al*. 2012; Vicente *et al*. 2022). *Lachancea thermotolerans* stands out for its ability to acidify wine through the production of lactic acid (up to 9 g/L) without significant increments in acetic acid under oenological conditions (Porter *et al*. 2019; Vicente *et al*. 2022; Vilela 2019). For that reason, it is one the most valuable yeasts used for acidity control, with some commercially available strains. Furthermore, *L. thermotolerans* improves wine aromatic complexity through the production of different aromas, e.g., 2-phenylethanol (Vicente *et al*. 2021a). For the isolation and selection of yeasts with these characteristics, the use of a simple, precise, and reproducible genotyping method is very convenient. Several attempts to study the *L. thermotolerans* intraspecific diversity have been accomplished. The first one described, analyzed the mitochondrial DNA of *L. thermotolerans* using restriction analysis approaches. This study showed a high homology among this species, without influence of the geographic or niche origin of strains (Belloch *et al*. 1997). Later, applying NGS techniques, the high conserved mitochondrial DNA structure was confirmed (Freel *et al*. 2014; Friedrich *et al*. 2012). Other approaches analyzing different microsatellites have been described to study of the intraspecific diversity of this species, being valuable for phylo-ecological studies (Banilas *et al*. 2016; Hranilovic *et al*. 2017). Nevertheless, these techniques are hardly implementable in winery-related strain selection procedures. The fluorescent labelled-multiplexed SSR analysis followed by a capillary electrophoresis revealed the influence of the geographical source of isolation in the population architecture in strains coming from several vineyards, a fact that was not supported applying other typing techniques based on tandem-repeat tRNA (Banilas *et al*. 2016) but later confirmed by deeper studies (Hranilovic *et al*. 2017).

With the specific objective of developing a comprehensive tool to search new *L. thermotolerans*, in this work a genotyping procedure for this species has been developed. This method can be applied to distinguish between *L. thermotolerans* strains in selection programs and to follow implantation of inoculated strains during winemaking or dry yeast production procedures. New primers have been designed to select those that allow a good resolution using agarose gel electrophoresis. Here we have verified the technique against several collections to prove its specificity and sensibility, both in collection and natural isolates. We tested the usefulness of the technique by following-up the implantation of an *in vitro* co-culture of *L. thermotolerans* strains under different complexity levels.

## 2. MATERIALS AND METHODS

### 2.1. Yeast strains and molecular identification of isolates

The yeast strains used as controls in this study, and coming from different environments and substrates, are listed in Table S1. Briefly, 103 *Lachancea thermotolerans* strains that were provided from different laboratories, culture collections and yeast producing companies were employed for technique verification. The specificity was assayed using several strains from other *Lachancea* species as well as other yeast genera. As well, a collection of *L. thermotolerans* autochthonous isolates coming from different fermentative stages (must, sulphited must and wines at 1040 and 999 densities) and from several vineyards of different Spanish wine appellations (Ribera de Duero, Rioja, and Manzanilla – Sanlúcar de Barrameda) in two consecutive vintages (2020 and 2021) is listed in Table S2. All yeasts strains were cryopreserved in 25% glycerol at −80°C. For yeast propagation, YMA agar plates were used (0.5% proteose peptone, 0.3% yeast extract, 0.3% malt extract, 1.0% glucose, 1.5% agar) and incubated at 28°C. Total genomic DNA was purified using the isopropanol method as described elsewhere (Querol *et al*. 1992) and stored at −20°C for further analysis.

### 2.2. Minisatellites identification and primers design

For microsatellite identification and primer design, the complete genome (including mitochondrial DNA) of the type strain *L. thermotolerans* CBS 6340 was downloaded from NCBI and used as template. Tandem repeated sequences were identified using the Tandem Repeats Finder described by Benson (Benson 1999) and studied as possible microsatellites candidates. Different loci by chromosome were selected according to the repeat length and the number of repeats. Primers (Table S3) were designed using Primer3 (https://primer3.ut.ee/) at the flanking regions of each minisatellite candidate. For specificity, primers were firstly tested using BLAST search and then *in vitro* using several strains.

### 2.3. Primer verification and selection for minisatellites fingerprinting amplification

Ten strains, representing a wide diversity in terms of geographic origin of the isolates, of *L. thermotolerans* (10-1488, CBS 10520, CBS 2907, CECT 1951, CONCERTO, DBVPG 3418, DMKU-RK 361, PYCC 4135, PYCC 6986 and UWOPS 85-312.1) were used in a first selection stage for the analysis of the most variable microsatellites. To select the primers that allowed the maximum discrimination between strains individual PCRs for each primer pair were performed in triplicate in a final volume of 25 μL containing 100 ng of genomic DNA, 2 μM of each primer (Integrated DNA Technologies, USA) and DreamTaq Green DNA polymerase 2x (ThermoFisher, USA). PCR was performed in a ProFlex PCR system (Applied Biosystems, USA) with an initial denaturation cycle at 95°C for 5 minutes, 25 cycles at 95°C for 1 minute, 55°C for 1 minute, and 72°C for 1.5 minutes and a final extension step at 72°C for 10 minutes. DNA electrophoresis was carried out using 15 μL of the PCR product in a 1.6 % (w/v) agarose gel and resolved in 1X TAE at 70 V for 110 minutes. After that, DNA was stained using a 1X GelRed solution (Biotium, USA) in 0.1 M NaCl. A 100 to 3,000 base pairs DNA weight marker (VWR, USA) was employed for band sizing. Gel images were captured employing a Gel Analyzer System (Axygen Scientific, USA).

### 2.4. Microsatellites fingerprinting and strain classification

The selected primers were multiplexed in a single PCR as described above with some modifications. The final concentration of B and F chromosome loci primers was reduced to 0.5 μM each. The rest of PCR conditions were as described above.

For strain classification, gel band analysis was performed using GelAnalyzer v.19.1 (www.gelanalyzer.com). Band fingerprinting of every gel was translated into binary (0 and 1) matrices. Clustering of isolates was done based on Sørensen-Dice coefficient and then represented by hierarchical clustering using the Ward calculation methods employing *ade4* package from R studio (Dray and Dufour 2007; R Core Team 2013). Correlation between genotypic and geographical distance matrices was calculated based on Mantel’s statistic based on Pearson’s product -moment correlation implemented in R studio, using *geosphere* and *vegan* packages.

### 2.5 Co-cultures for strain monitorization

To assess the adequacy of the technique to monitorization strain implantation during wine fermentations, different co-culture assays were developed. Four strains (ROD21-99, A11-606, A11-612 and UWOPS 79-116) were selected to perform the *in-vitro* co-cultures at different complexity levels: two-strain, three-strain, and four-strain communities. All the cultures were inoculated at an initial cellular density of 10^6^ cells/mL in 2.5 mL using 12-well microtiter plates. 100 μL samples were taken at 0, 24 and 96 hours, serially diluted and spread on YMA plates. From each sample, 20 colonies per community were randomly taken, genotyped according to the hereto described technique and compared to the fingerprint of the strains in pure culture. Then, implantation percentages were calculated for each strain in each community and time

## 4. RESULTS AND DISCUSSION

Here we present an accurate genotyping method for *L. thermotolerans* selection and strain monitorization. The genome, including mitochondrial DNA, of the type strain of *L*. *thermotolerans* CBS 6340 was analysed for the detection of microsatellites. In total, 1,038 Simple Sequence Repeats (SSR) were present in this reference genome, 23 of them located in the mitochondrial chromosome. SSR were filtered according to the repetition length and the predicted number on repetitions, maintaining only the longest and those presented in high copy number. Finally, the most adequate microsatellite candidates were selected for *in vitro* testing. The selected microsatellites and its primers for PCR are described in Table S3. The first approach revealed the unsuitability of the mitochondrial microsatellites since all the ten strains that were initially used presented the same amplification products according to that previously reported (Belloch *et al*. 1997; Freel *et al*. 2014; Friedrich *et al*. 2012).

The genomic-located loci showed a better performance, nevertheless, some of them indicated a great homogeneity among the studied strains. In some cases, no amplification was detected for different primers in several strains. Despite this fact, those primers were not discarded since they allow a differentiation among strains in a presence/absence criterion. At the end, six of them, located in different chromosomes (A, B, F, G, and H) were selected by two reasons. They were those that presented the higher divergence among all the strains tested and, secondly, the combination of twelve different primers is suitable enough for a multiplexed PCR.

The analysis, that was performed using five different *Lachancea non-thermotolerans* species and thirty-four yeasts isolates belonging to genera different from *Lachancea*, showed a great specificity. No amplification products were obtained in any case, indicating that this is a species-specific tool. This fact allows a direct isolate-fingerprinting analysis without requiring a previously molecular (e.g., 26S rDNA sequencing) identification of the isolate as occur with the interdelta genotyping method (Legras & Karst, 2003). The application of this technique in more than one hundred collection and reliable strains reached excellent results, allowing the differentiation of 99 out of the 103 analysed strains, which means a 96 % of sensibility (Figure 1) (Table S4). The genotypic clustering obtained by employing these microsatellites lacked geographical significance since the genotypic differences of the strains were not related with the geographical origin of the isolate, as Mantel correlation test showed (p=0.658) (Figure S1-A).

**Figure 1.**
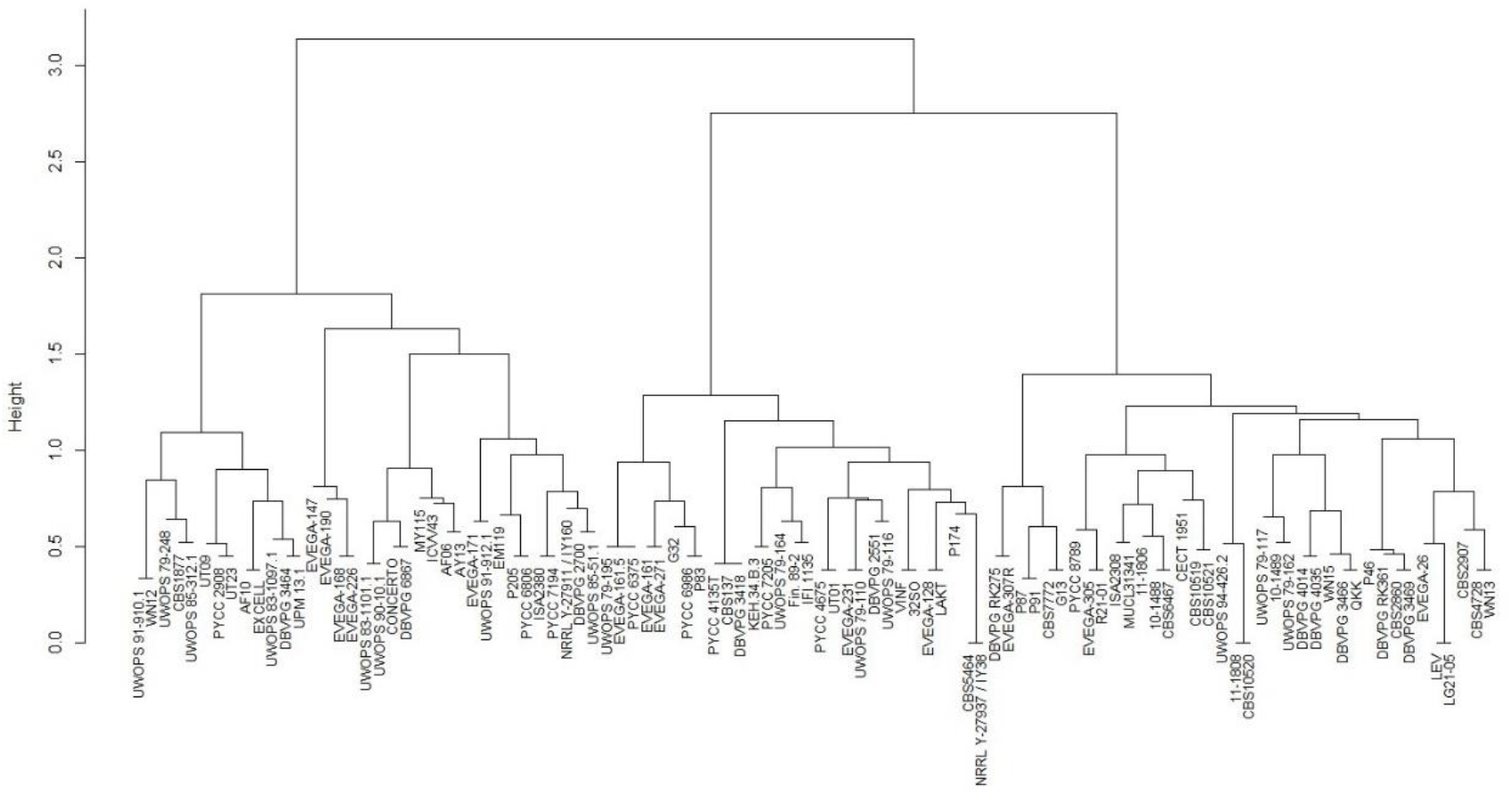
Dendrogram showing the different clusters based on Sørensen-Dice coefficient constructed using Ward’s methods for *L. thermotolerans* collection strains.

The application of the procedure in a collection of autochthonous isolates coming from several vineyards and harvests allowed the differentiation of 190 different fingerprints among the 458 isolates studied, which means that around the 45 % of the isolates presented a unique fingerprint (Figure S2, Table S5). The diversity present among the *L. thermotolerans* isolates was extremely high. These results are similar to those showed by others studying the intraspecific diversity of *L. thermotolerans* in Greek isolates, that is extremely high (Banilas *et al*. 2016). Nevertheless, this data did not show any kind of population structure since any geographical clustering significance was observed (p=0.03) (Figure S1-B).

In our study, it was not possible to discriminate the strains by their geographical distribution, unlike the results obtained in Banilas’ study, where the strains were isolated from two winemaking regions separated by about 500 km and the sea in between, that conforms an important geographical barrier for allopatric differentiation. The increase in the number of microsatellites may improve its significance for geographical dispersion studies, since when we reduced the geographical dispersion of our isolates, the significance level was much lower (isolated strains compared to collection ones). As well, the diversity impact in wine fermentation is still partially unknown for *S. cerevisiae* as well as for other yeast species. Some *non-Saccharomyces* species, such as *Hanseniaspora uvarum* and *Starmerella bacillaris*, show a great diversity (Masneuf-Pomarede *et al*. 2016) as *L. thermotolerans*. This diversity has been probably driven by the selective pressure in wine-related environments (Hranilovic *et al*. 2017). The results concerning the *terroir* designation (regarding geographical influence) in non-*Saccharomyces* species is unclear. Some genetic patterns regarding *S. cerevisiae* have been described, confirming the singularity of some stains in a certain geographical location (de Celis *et al*. 2019), fact that has not been confirmed in other wine-related species (Banilas *et al*. 2016). Despite this fact, the genetic profile is not the unique condition that confirms the uniqueness of a strain, the oenological phenotypes are essential for *terroir* confirmation.

Finally, we carried out an additional verification of the PCR technique to test the suitability for strain monitoring studies, both in wine fermentation and dry yeast biomass production. With this purpose we performed several co-cultures containing different strains of *L. thermotolerans* mixed up at different complexity levels and different times, from 0, to check the initial inoculation ratio, to 96 hours, when the lactic acid production peaked, and *S. cerevisiae* is usually inoculated in sequential fermentations involving both species. The application of the technique for wine-monitoring purposes showed the accuracy of the PCR multiplex-based genotyping method hereto presented. The evolution of the strains expressed as implantation percentages are shown in Table 1. We were able to track every strain along the fermentation since each strain showed its characteristic band profile in all the analysis. Huge differences were observed among the implantation capacity of the strains. Some strains (e.g., UWOPS 79-116) were able to grow faster in the conditions tested, displacing others, and becoming dominant at the end of the fermentation trials. This fact is of great importance since, in rational strain selection procedures, one of the most valuable characteristics is the rapid growth and dominance of the selected strain over the autochthonous microbiota. This strain selection procedures will be essential for building synthetic yeast starter-culture consortia with microbial *terroir* effects (Pretorius 2020).

**Table 1.** Implantation percentages of each strain used in the synthetic *L. thermotolerans* communities at different sampling times.

| Community | Sampling time (h) | Incidence (%) |  |  |  |
| --- | --- | --- | --- | --- | --- |
|  |  | A11-606 | ROD21-99 | UWOPS 79-116 | A11-612 |
| A | 0 | 55 | 45 | - | - |
|  | 24 | 35 | 65 | - | - |
|  | 96 | 5 | 95 | - | - |
| B | 0 | 50 | 30 | 20 | - |
|  | 24 | 40 | 45 | 15 | - |
|  | 96 | 0 | 40 | 60 | - |
| C | 0 | 25 | 25 | 25 | 25 |
|  | 24 | 0 | 25 | 20 | 55 |
|  | 96 | 0 | 35 | 40 | 25 |

The simplicity, reproducibility, and valuable results that this technique achieves are extremely applicable in wine-related strain selection procedures. The method has been tested in different groups of strains, both from collection and natural origin. The first trial allowed us to determine the discrimination capacity of our method; the second one tested the real performance of it in a strain selection procedure. The final validation showed valuable results since every strain can be followed-up along the time even in high-complexity *L. thermotolerans* communities. So, the method here described is a simple and implementable procedure to allow, not only the strain classification, but the fermentation monitoring as well as the strain competition capacity that defines it performance in different industrial applications.

## 5. ACKNOWLEDGMENTS

Funding for the research in this paper was provided by the Spanish Ministry of Science and Innovation under the framework of the CDTI project LowpHWine (IDI-20210391) and project VinoSegCalClim (PID2020-119008RB-I00). We thank Marc-André Lachance (Western University, Canada), Joseph Schacherer (University of Strasbourg, France), Matthias Sipiczki (University of Debrecen, Hungary), Manuel Malfeito (University of Lisbon, Portugal), Pilar Blanco (Galician Viticulture and Enology Station, Spain), Ana Rosa Gutiérrez (Institute of Vine and Wine Sciences, Spain), Pilar Santamaría (Institute of Vine and Wine Sciences, Spain) and María Victoria Moreno-Arribas (Food Science Research Institute, Spain) for the kindly transfer of several yeast strains.

**Table Supplementary 1.** Collection strains used in this study as positive and negative controls. Strain code, identification, original collection, geographical and isolation source.

**Table Supplementary 2.** Wine related *L. thermotolerans* isolates used in this study.

Isolate code, identification, fermentative stage, winery, and harvest.

**Table Supplementary 3.** Designed primers of the study for multiplex-PCR. Chromosome location, primer name, sequence and melting temperature. In bold, those employed for multilocus typing of *L. thermotolerans*.

**Table Supplementary 4.** Amplicon size for each *L. thermotolerans* collection strain used in this study as positive controls. Strain code and molecular size determined using GelAnalyzer (there are as many entries for a single isolate as the number of amplicons present).

**Table Supplementary 5.** Amplicon size for each *L. thermotolerans* wine related isolates used in this study. Strain code and molecular size determined using GelAnalyzer (there are as many entries for a single isolate as the number of amplicons present).

**Figure Supplementary 1.** Dendrogram showing the different clusters based on Sørensen-Dice coefficient constructed using Ward’s methods for *L. thermotolerans* natural isolates.

