## Supplementary material for "Multilocus typing of *Lachancea thermotolerans* for wine fermentation monitoring"

[1]Department of Genetics, Physiology and Microbiology. Unit of Microbiology. Biology Faculty, Complutense University of Madrid, 28040 Madrid, Spain

[2]Pago de Carraovejas, S.L.U., Camino de Carraovejas, s/n, 47300 Peñafiel, Valladolid, Spain

[3]Department of Chemistry and Food Technology, Polytechnic University of Madrid, Ciudad Universitaria s/n, 28040 Madrid, Spain

**Table Supplementary 1.** Collection strains used in this study as positive and negative controls. Strain code, identification, original collection, geographical and isolation source.

| STRAIN CODE | IDENTIFICATION | OTHER COLLECTIONS | ORIGIN | GEOGRAPHYCAL SOURCE | ISOLATION SOURCE |
|---|---|---|---|---|---|
| 10-1488 | Lachancea thermotolerans | CCY 043-013-005 | M. Sipiczki, Debrecen University, Hungary | Černochov; Slovakia | Mummified grapes |
| 10-1489 | Lachancea thermotolerans | CCY 043-013-004 | M. Sipiczki, Debrecen University, Hungary | Černochov; Slovakia | Mummified grapes |
| 11-1806 | Lachancea thermotolerans | | M. Sipiczki, Debrecen University, Hungary | | |
| 11-1808 | Lachancea thermotolerans | | M. Sipiczki, Debrecen University, Hungary | | |
| CBS10519 | Lachancea thermotolerans | YJS4203 | Vang - INRA Grignon | Ukraine | exsudate of Quercus sp. |
| CBS10520 | Lachancea thermotolerans | YJS4204 | Vang - INRA Grignon | Russia | exsudate of Quercus mongolica |
| CBS10521 | Lachancea thermotolerans | YJS4205 | Vang - INRA Grignon | Finland | exsudate of Quercus sp. |
| CBS137 | Lachancea thermotolerans | YJS4206 | Vang - INRA Grignon | Netherlands | date |
| CBS1877 | Lachancea thermotolerans | YJS4207 | Vang - INRA Grignon | Italy | grape |
| CBS2860 | Lachancea thermotolerans | YJS4209 | Vang - INRA Grignon | Sardinia | grape must |
| CBS2907 | Lachancea thermotolerans | YJS4210 | Vang - INRA Grignon | South Africa | soil |
| CBS4728 | Lachancea thermotolerans | YJS4212 | Vang - INRA Grignon | Czechoslovakia | grape |
| CBS5464 | Lachancea thermotolerans | YJS4213 | Vang - INRA Grignon | Australia | cotton seed |
| CBS6467 | Lachancea thermotolerans | YJS4218 | Vang - INRA Grignon | Japan | exudate of tree |
| CBS7772 | Lachancea thermotolerans | YJS4219 | Vang - INRA Grignon | Brazil | Uca sp. (fiddler crab) |
| CECT 1951 | Lachancea thermotolerans | IFO1050 , NCYC 412 | Spanish Culture Collection | | |
| Concerto | Lachancea thermotolerans | | CHR-Hansen, Denmark | Mediterranean country | |
| DBVPG 2551 | Lachancea thermotolerans | YJS4254 | DBVPG - Italy collection | Piemonte, Italy | barbera wine |
| DBVPG 2700 | Lachancea thermotolerans | YJS4255 | DBVPG - Italy collection | Campo de Santiago, La Mancha, Spain | airen grapes |
| DBVPG 3418 | Lachancea thermotolerans | YJS4256 | DBVPG - Italy collection | Italy | milk |
| DBVPG 3464 | Lachancea thermotolerans | YJS4257 | DBVPG - Italy collection | ValdepeÑas, La Mancha, Spain | grapes |
| DBVPG 3466 | Lachancea thermotolerans | YJS4258 | DBVPG - Italy collection | La Encomienda, La Mancha, Spain | grapes |
| DBVPG 3469 | Lachancea thermotolerans | YJS4259 | DBVPG - Italy collection | Manzanares, La Mancha, Spain | grapes |
| DBVPG 4014 | Lachancea thermotolerans | YJS4260 | DBVPG - Italy collection | Italy | caverns |
| DBVPG 4035 | Lachancea thermotolerans | YJS4261 | DBVPG - Italy collection | ex Yugoslavia | grape must |
| DBVPG 6867 | Lachancea thermotolerans | YJS4262 | DBVPG - Italy collection | Brazil | Pilosocereus arrabidae, cactus |
| DMKU-RK275 | Lachancea thermotolerans | YJS4248 | Savitree | Thailand | rice leaves |
| DMKU-RK361 | Lachancea thermotolerans | YJS4249 | Savitree | Nong Muang, Thailand | sugarcane leaves |
| Fin. 89-2 | Lachancea thermotolerans | YJS4263 | Naumova | Finland | exudate of Quercus sp. |
| KEH.34.B.3 | Lachancea thermotolerans | YJS4247 | Justin Fay | Ste. Genevieve, Missouri, USA | Chaumette vineyard fermentation, fermentation |
| MUCL31341 | Lachancea thermotolerans | YJS4222 | Vang - INRA Grignon | Italy | wine |
| NRRL Y-27911 / IY160 | Lachancea thermotolerans | YJS4252 | Meredith Blackwell | LA, USA | gut of male Chauliodes rastricornis |
| NRRL Y-27937 / IY38 | Lachancea thermotolerans | YJS4251 | Meredith Blackwell | LA, USA | surface of male Chauliodes rastricornis |
| PYCC 2908 | Lachancea thermotolerans | CBS 2917 | Portuguese Yeast Culture Collection | | Drosophila sp |
| PYCC 4135T | Lachancea thermotolerans | CBS 6340$^{T}$ | Portuguese Yeast Culture Collection | Russia/Asia | mirabelle plum conserve |
| PYCC 4675 | Lachancea thermotolerans | | Portuguese Yeast Culture Collection | Oeiras, Portugal | |
| PYCC 6375 | Lachancea thermotolerans | | Portuguese Yeast Culture Collection | Arteixo, Galicia, España | Soil under Q pirenaica |
| PYCC 6806 | Lachancea thermotolerans | OR967 | Portuguese Yeast Culture Collection | Arrabida Park, Portugal | Soil |
| PYCC 6986 | Lachancea thermotolerans | | Portuguese Yeast Culture Collection | Evolution canyon, Israel | Tree hole with sap leaking from Ceratonia siliqua |
| PYCC 7194 | Lachancea thermotolerans | | Portuguese Yeast Culture Collection | Santarem, Portugal | Fallen leaves from olive trees |
| PYCC 7205 | Lachancea thermotolerans | | Portuguese Yeast Culture Collection | Alqueva, Portugal | Olives from tree |
| PYCC 8789 | Lachancea thermotolerans | | Portuguese Yeast Culture Collection | Alqueva, Portugal | Olives from tree |
| UWOPS 79-110 | Lachancea thermotolerans | | M. A. Lachance, Western University, Ontario, Canada | Ontario, Canada | black knot, Prunus virginiana |
| UWOPS 79-116 | Lachancea thermotolerans | | M. A. Lachance, Western University, Ontario, Canada | Pinery, Canada | black knot, Prunus virginiana |
| UWOPS 79-117 | Lachancea thermotolerans | | M. A. Lachance, Western University, Ontario, Canada | Pinery, Canada | black knot, Prunus virginiana |
| UWOPS 79-162 | Lachancea thermotolerans | | M. A. Lachance, Western University, Ontario, Canada | Melbourne, Ontario, Canada | black knot, Prunus virginiana |
| UWOPS 79-164 | Lachancea thermotolerans | | M. A. Lachance, Western University, Ontario, Canada | Melbourne, Ontario, Canada | black knot, Prunus virginiana |
| UWOPS 79-195 | Lachancea thermotolerans | | M. A. Lachance, Western University, Ontario, Canada | Melbourne, Ontario, Canada | black knot, Prunus virginiana |
| UWOPS 79-248 | Lachancea thermotolerans | | M. A. Lachance, Western University, Ontario, Canada | Ontario, Canada | frass, Birch |
| UWOPS 83-1097.1 | Lachancea thermotolerans | | M. A. Lachance, Western University, Ontario, Canada | Cayman Brac, Cayman Islands | Gitonia americana, Opuntia Stricta |
| UWOPS 83-1101.1 | Lachancea thermotolerans | | M. A. Lachance, Western University, Ontario, Canada | Cayman Brac, Cayman Islands | Gitonia americana, Opuntia Stricta |
| UWOPS 85-312.1 | Lachancea thermotolerans | | M. A. Lachance, Western University, Ontario, Canada | Tuscon, Arizona, USA | Drosophila carbonaria, Propis juliflora |
| UWOPS 85-51.1 | Lachancea thermotolerans | | M. A. Lachance, Western University, Ontario, Canada | Big Pine Key, Florida, USA | Opuntia cubensis |
| UWOPS 90-10.1 | Lachancea thermotolerans | | M. A. Lachance, Western University, Ontario, Canada | Shroud Cay, Exumas Cay, Bahamas | Columnar cactus |
| UWOPS 91-910.1 | Lachancea thermotolerans | | M. A. Lachance, Western University, Ontario, Canada | Saddle Rd Park, Hawaii | flux (pink), Myoporum |
| UWOPS 91-912.1 | Lachancea thermotolerans | | M. A. Lachance, Western University, Ontario, Canada | Saddle Rd Park, Hawaii | flux (pink), Myoporum |
| UWOPS 94-426.2 | Lachancea thermotolerans | | M. A. Lachance, Western University, Ontario, Canada | Jalisco, Mexico | destilery, agave must |
| ISA 2380 | Lachancea thermotolerans | Pi4 | M. Malfeito, Universidad de Minho, Portugal | | sour rot must |
| ISA 2308 | Lachancea thermotolerans | 44c | M. Malfeito, Universidad de Minho, Portugal | | |
| Excellance | Lachancea thermotolerans | | Lamothe-Abiet, France | | |
| Laktia | Lachancea thermotolerans | | Lallemand, Canada | | |
| Levulia | Lachancea thermotolerans | | AEB, Italy | | |
| EnartisFerm Qk | Lachancea thermotolerans | | Enartis, Italy | | |
| Viniflora | Lachancea thermotolerans | | CHR-Hansen, Denmark | | |
| IFI 1135 | Lachancea thermotolerans | CECT 12813 | Food Science Research Institute, Superior Council of Scientific Investigations, Spain | Solana, Spain | Airen grapes |
| UPM 13.1 | Lachancea thermotolerans | | Polytechnic University of Madrid | | |
| 32SO | Lachancea thermotolerans | | Food Science Research Institute, Superior Council of Scientific Investigations, Spain | | |
| EVEGA-128 | Lachancea thermotolerans | | EVEGA, Spain | DO Monterei, Galicia, Spain | mencia grape, ecologic vineyard |
| EVEGA-147 | Lachancea thermotolerans | | EVEGA, Spain | DO Ribeiro, Galicia, Spain | ecologic vineyard |
| EVEGA-231 | Lachancea thermotolerans | | EVEGA | DO Monterei, Galicia, Spain | mencia grape, ecologic vineyard |
| EVEGA-26 | Lachancea thermotolerans | | EVEGA | DO Riberira Sacra, Galicia, Spain | mencia grape, conventional vineyard |
| EVEGA-271 | Lachancea thermotolerans | | EVEGA | DO Ribeiro, Galicia, Spain | treixadura grape, conventional vineyard |
| EVEGA-305 | Lachancea thermotolerans | | EVEGA | DO Ribeiro, Galicia, Spain | treixadura grape, conventional vineyard |
| EVEGA-307R | Lachancea thermotolerans | | EVEGA | DO Ribeiro, Galicia, Spain | treixadura grape, conventional vineyard |
| EVEGA-171 | Lachancea thermotolerans | | EVEGA | DO Ribeiro, Galicia, Spain | Brancellao grape, ecologic vineyard |
| EVEGA-168 | Lachancea thermotolerans | | EVEGA | DO Ribeiro, Galicia, Spain | Brancellao grape, conventional vineyard |
| EVEGA-161.5 | Lachancea thermotolerans | | EVEGA | DO Ribeiro, Galicia, Spain | treixadura grape, ecologic vineyard |
| EVEGA-161 | Lachancea thermotolerans | | EVEGA, Spain | DO Ribeiro, Galicia, Spain | Brancellao grape, conventional vineyard |
| EVEGA-190 | Lachancea thermotolerans | | EVEGA, Spain | DO Ribeiro, Galicia, Spain | treixadura grape, conventional vineyard |
| EVEGA-226 | Lachancea thermotolerans | | EVEGA, Spain | DO Monterei, Galicia, Spain | mencia grape, ecologic vineyard |
| AY13 | Lachancea thermotolerans | | ICVV, Spain | Aldeanueva de Ebro, Spain | |
| ICVV43 | Lachancea thermotolerans | | ICVV, Spain | Samaniego, Spain | |
| MY115 | Lachancea thermotolerans | | ICVV, Spain | Mendavia, Sapin | |
| UT01 | Lachancea thermotolerans | | Complutense Yeast Collection | Utiel, Spain | bobal grapes, ecologic vineyard, 2020 |
| UT23 | Lachancea thermotolerans | | Complutense Yeast Collection | Utiel, Spain | bobal grapes, ecologic vineyard, 2020 |
| UT9 | Lachancea thermotolerans | | Complutense Yeast Collection | Utiel, Spain | bobal grapes, ecologic vineyard, 2020 |
| AF06 | Lachancea thermotolerans | | Complutense Yeast Collection | Alfaro, Spain | tempranillo grapes, conventional vineyard 2020 |
| AF10 | Lachancea thermotolerans | | Complutense Yeast Collection | Alfaro, Spain | tempranillo grapes, conventional vineyard 2020 |
| WN12 | Lachancea thermotolerans | | Complutense Yeast Collection | Salamanca, Spain | thyme flower, 2021 |
| WN13 | Lachancea thermotolerans | | Complutense Yeast Collection | Salamanca, Spain | clover flower, 2021 |
| WN15 | Lachancea thermotolerans | | Complutense Yeast Collection | Salamanca, Spain | soil under quercus ilex, 2021 |
| G13 | Lachancea thermotolerans | | Complutense Yeast Collection | Leon, Spain | viura grapes, 2012 |
| G32 | Lachancea thermotolerans | | Complutense Yeast Collection | Leon, Spain | viura grapes, 2012 |
| P83 | Lachancea thermotolerans | | Complutense Yeast Collection | Peñafiel, Spain | tempranillo grapes, 2013 |
| P91 | Lachancea thermotolerans | | Complutense Yeast Collection | Peñafiel, Spain | tempranillo grapes, 2013 |
| P174 | Lachancea thermotolerans | | Complutense Yeast Collection | Peñafiel, Spain | tempranillo grapes, 2014 |
| P205 | Lachancea thermotolerans | | Complutense Yeast Collection | Peñafiel, Spain | tempranillo grapes, 2014 |
| P46 | Lachancea thermotolerans | | Complutense Yeast Collection | Peñafiel, Spain | tempranillo grapes, 2013 |
| P87 | Lachancea thermotolerans | | Complutense Yeast Collection | Peñafiel, Spain | tempranillo grapes, 2013 |
| EM119 | Lachancea thermotolerans | | Complutense Yeast Collection | Pesquera de Duero, Spain | 2014 |
| LG21-05 | Lachancea thermotolerans | | Complutense Yeast Collection | Languedoc, France | cariñena grape, 2021 |
| R21-01 | Lachancea thermotolerans | | Complutense Yeast Collection | Calahorra, Spain | tempranillo grape, 2021 |
| CECT 11056 | Lachancea fermentati | CBS 707, NRRL Y-1559T | Spanish Culture Collection | | |
| CECT 1965 | Lachancea waltii | CBS 6430T | Spanish Culture Collection | | |
| PYCC 2817 | Lachancea kluyveri | CBS 3082T | Portuguese Yeast Culture Collection | | |
| PYCC 6985 | Lachancea cidri | ZP 995 | Portuguese Yeast Culture Collection | | |
| PYCC 6533T | Lachancea lanzarotensis | CBS 12615T;CECT 13066 | Portuguese Yeast Culture Collection | | |
| Ap_NS-O-70 | Aureobasidium pullulans | | Complutense Yeast Collection | | |
| Ba_OGB1 | Brettanomyces anomalus | | Complutense Yeast Collection | | |
| Bb_5B | Brettanomyces bruxelensis | | Complutense Yeast Collection | | |
| Cz_NS-WN-007 | Candida zemplinina | | Complutense Yeast Collection | | |
| Co_CR-111 | Candida oleophila | | Complutense Yeast Collection | | |
| Ct_UT-024 | Candida tropicalis | | Complutense Yeast Collection | | |
| Cf_LR-001 | Cyberlindmera fabianii | | Complutense Yeast Collection | | |
| Ca_NS-PDC-132 | Cryptococcus amylolentus | | Complutense Yeast Collection | | |
| De_NS-WN-006 | Debaryomyces sp. | | Complutense Yeast Collection | | |
| Hos_DN-411 | Hanseniaspora osmophila | | Complutense Yeast Collection | | |
| Hop_BM-001 | Hanseniaspora opuntiae | | Complutense Yeast Collection | | |
| Hu_NS-O-4 | Hanseniaspora uvarum | | Complutense Yeast Collection | | |
| Ku_DN-201 | Kazachtania unispora | | Complutense Yeast Collection | | |
| Km_RDG-003 | Kluyveromyces marxianus | | Complutense Yeast Collection | | |
| Km_NS-PDC-100 | Kluyveromyces marxianus | | Complutense Yeast Collection | | |
| Le_MR-204 | Lodderomyces elongisporus | | Complutense Yeast Collection | | |
| Le_BKM12.5 | Lodderomyces elongisporus | | Complutense Yeast Collection | | |
| Mp_ME-004 | Metchsnikowia pulcherrima | | Complutense Yeast Collection | | |
| Mf_NS-O-29 | Metchsnikowia fructicola | | Complutense Yeast Collection | | |

| | | | |
|---|---|---|---|
| Mv_NS-O-32 | Metchsnikowia viticola | | Complutense Yeast Collection |
| Mg_NS-G-57 | Meyerozyma guillermondii | | Complutense Yeast Collection |
| Ni_BKM17.1 | Nakazawaea ishiwadae | | Complutense Yeast Collection |
| Ng_CR-112 | Naganishia globosa | | Complutense Yeast Collection |
| Pm_MR-412 | Pichia membranifaciens | | Complutense Yeast Collection |
| Pk_MR-321 | Pichia kudriavzevii | | Complutense Yeast Collection |
| Rg_NS-G-61 | Rhodotorula glutinis | | Complutense Yeast Collection |
| Rm_LS-108 | Rhodotorula mucilaginosa | | Complutense Yeast Collection |
| Sc_AG012 | Saccharomyces cerevisiae | | Complutense Yeast Collection |
| Sc_AG023 | Saccharomyces cerevisiae | | Complutense Yeast Collection |
| Sp_UPMSP-936 | Schizosaccharomyces pombe | | Complutense Yeast Collection |
| Td_NS-TD | Torulaspora delbrueckii | | Complutense Yeast Collection |
| Wa_NS-G-034 | Wickerhamomyces anomalus | | Complutense Yeast Collection |
| Zb-NS-G-063 | Zygosaccharomyces bailii | | Complutense Yeast Collection |
| Pm-MR-339 | Pichia manshurica | | Complutense Yeast Collection |

**Table Supplementary 2.** Wine related *L. thermotolerans* isolates used in this study. Isolate code, identification, fermentative stage, winery, and harvest.

| Isolate | Identification | Fermentative stage | Winery | Harvest |
|---|---|---|---|---|
| A11-603 | *Lachancea thermotolerans* | 2. T24 | P | 2021 |
| A11-606 | *Lachancea thermotolerans* | 2. T24 | P | 2021 |
| A11-612 | *Lachancea thermotolerans* | 2. T24 | P | 2021 |
| BAR20-2.2 | *Lachancea thermotolerans* | n.d. | B | 2020 |
| BAR20-2.39 | *Lachancea thermotolerans* | n.d. | B | 2020 |
| BAR21-1.11 | *Lachancea thermotolerans* | n.d. | B | 2021 |
| BAR21-1.12 | *Lachancea thermotolerans* | n.d. | B | 2021 |
| BAR21-1.13 | *Lachancea thermotolerans* | n.d. | B | 2021 |
| BAR21-1.14 | *Lachancea thermotolerans* | n.d. | B | 2021 |
| BAR21-1.15 | *Lachancea thermotolerans* | n.d. | B | 2021 |
| BAR21-1.18 | *Lachancea thermotolerans* | n.d. | B | 2021 |
| BAR21-1.19 | *Lachancea thermotolerans* | n.d. | B | 2021 |
| BAR21-1.20 | *Lachancea thermotolerans* | n.d. | B | 2021 |
| BAR21-1.4 | *Lachancea thermotolerans* | n.d. | B | 2021 |
| BAR21-1.6 | *Lachancea thermotolerans* | n.d. | B | 2021 |
| BD-522 | *Lachancea thermolerans* | 1. T0 | H | 2021 |
| BD-525 | *Lachancea thermolerans* | 1. T0 | H | 2021 |
| BD-601 | *Lachancea thermolerans* | 2. T24 | H | 2021 |
| BD-612 | *Lachancea thermolerans* | 2. T24 | H | 2021 |
| BD-619 | *Lachancea thermolerans* | 2. T24 | H | 2021 |
| BD-705 | *Lachancea thermolerans* | 3. D1040 | H | 2021 |
| BD-707 | *Lachancea thermolerans* | 3. D1040 | H | 2021 |
| BD-714 | *Lachancea thermolerans* | 3. D1040 | H | 2021 |
| BD-715 | *Lachancea thermolerans* | 3. D1040 | H | 2021 |
| CB-1-01 | *Lachancea thermotolerans* | 1. T0 | P | 2020 |
| CB-1-03 | *Lachancea thermotolerans* | 1. T0 | P | 2020 |
| CB-1-07 | *Lachancea thermotolerans* | 1. T0 | P | 2020 |
| CB-1-08 | *Lachancea thermotolerans* | 1. T0 | P | 2020 |
| CB-1-09 | *Lachancea thermotolerans* | 1. T0 | P | 2020 |
| CB-1-11 | *Lachancea thermotolerans* | 1. T0 | P | 2020 |
| CB-1-12 | *Lachancea thermotolerans* | 1. T0 | P | 2020 |
| CB-1-15 | *Lachancea thermotolerans* | 1. T0 | P | 2020 |
| CB-1-16 | *Lachancea thermotolerans* | 1. T0 | P | 2020 |
| CB-2-01 | *Lachancea thermotolerans* | 2. T24 | P | 2020 |
| CB-2-02 | *Lachancea thermotolerans* | 2. T24 | P | 2020 |
| CB-2-04 | *Lachancea thermotolerans* | 2. T24 | P | 2020 |
| CB-2-05 | *Lachancea thermotolerans* | 2. T24 | P | 2020 |
| CB-2-06 | *Lachancea thermotolerans* | 2. T24 | P | 2020 |
| CB-2-16 | *Lachancea thermotolerans* | 2. T24 | P | 2020 |
| CB-2-17 | *Lachancea thermotolerans* | 2. T24 | P | 2020 |
| CB-3-11 | *Lachancea thermotolerans* | 3. D1040 | P | 2020 |
| CB-3-12 | *Lachancea thermotolerans* | 3. D1040 | P | 2020 |
| CB-3-13 | *Lachancea thermotolerans* | 3. D1040 | P | 2020 |
| CB-3-15 | *Lachancea thermotolerans* | 3. D1040 | P | 2020 |
| CB-3-16 | *Lachancea thermotolerans* | 3. D1040 | P | 2020 |
| CB-3-17 | *Lachancea thermotolerans* | 3. D1040 | P | 2020 |
| CB-3-19 | *Lachancea thermotolerans* | 3. D1040 | P | 2020 |
| CB-3-20 | *Lachancea thermotolerans* | 3. D1040 | P | 2020 |
| CB-3-21 | *Lachancea thermotolerans* | 3. D1040 | P | 2020 |

| | | | | |
|---|---|---|---|---|
| CB-501 | *Lachancea thermotolerans* | 1. T0 | P | 2021 |
| CB-502 | *Lachancea thermotolerans* | 1. T0 | P | 2021 |
| CB-503 | *Lachancea thermotolerans* | 1. T0 | P | 2021 |
| CB-504 | *Lachancea thermotolerans* | 1. T0 | P | 2021 |
| CB-505 | *Lachancea thermotolerans* | 1. T0 | P | 2021 |
| CB-506 | *Lachancea thermotolerans* | 1. T0 | P | 2021 |
| CB-507 | *Lachancea thermotolerans* | 1. T0 | P | 2021 |
| CB-508 | *Lachancea thermotolerans* | 1. T0 | P | 2021 |
| CB-509 | *Lachancea thermotolerans* | 1. T0 | P | 2021 |
| CB-515 | *Lachancea thermotolerans* | 1. T0 | P | 2021 |
| CB-516 | *Lachancea thermotolerans* | 1. T0 | P | 2021 |
| CB-519 | *Lachancea thermotolerans* | 1. T0 | P | 2021 |
| CB-520 | *Lachancea thermotolerans* | 1. T0 | P | 2021 |
| CB-522 | *Lachancea thermotolerans* | 1. T0 | P | 2021 |
| CB-524 | *Lachancea thermotolerans* | 1. T0 | P | 2021 |
| CB-601 | *Lachancea thermotolerans* | 2. T24 | P | 2021 |
| CB-602 | *Lachancea thermotolerans* | 2. T24 | P | 2021 |
| CB-603 | *Lachancea thermotolerans* | 2. T24 | P | 2021 |
| CB-613 | *Lachancea thermotolerans* | 2. T24 | P | 2021 |
| CB-616 | *Lachancea thermotolerans* | 2. T24 | P | 2021 |
| CB-618 | *Lachancea thermotolerans* | 2. T24 | P | 2021 |
| CB-705 | *Lachancea thermotolerans* | 3. D1040 | P | 2021 |
| CB-708 | *Lachancea thermotolerans* | 3. D1040 | P | 2021 |
| CB-710 | *Lachancea thermotolerans* | 3. D1040 | P | 2021 |
| CB-713 | *Lachancea thermotolerans* | 3. D1040 | P | 2021 |
| CB-714 | *Lachancea thermotolerans* | 3. D1040 | P | 2021 |
| CB-716 | *Lachancea thermotolerans* | 3. D1040 | P | 2021 |
| CB-719 | *Lachancea thermotolerans* | 3. D1040 | P | 2021 |
| CB-801 | *Lachancea thermotolerans* | 4. D999 | P | 2021 |
| CR-1-01 | *Lachancea thermotolerans* | 1. T0 | H | 2020 |
| CR-1-03 | *Lachancea thermotolerans* | 1. T0 | H | 2020 |
| CR-1-04 | *Lachancea thermotolerans* | 1. T0 | H | 2020 |
| CR-1-06 | *Lachancea thermotolerans* | 1. T0 | H | 2020 |
| CR-1-07 | *Lachancea thermotolerans* | 1. T0 | H | 2020 |
| CR-1-08 | *Lachancea thermotolerans* | 1. T0 | H | 2020 |
| CR-1-09 | *Lachancea thermotolerans* | 1. T0 | H | 2020 |
| CR-1-13 | *Lachancea thermotolerans* | 1. T0 | H | 2020 |
| CR-1-15 | *Lachancea thermotolerans* | 1. T0 | H | 2020 |
| CR-1-16 | *Lachancea thermotolerans* | 1. T0 | H | 2020 |
| CR-1-20 | *Lachancea thermotolerans* | 1. T0 | H | 2020 |
| CR-2-02 | *Lachancea thermotolerans* | 2. T24 | H | 2020 |
| CR-2-03 | *Lachancea thermotolerans* | 2. T24 | H | 2020 |
| CR-2-04 | *Lachancea thermotolerans* | 2. T24 | H | 2020 |
| CR-2-10 | *Lachancea thermotolerans* | 2. T24 | H | 2020 |
| CR-2-11 | *Lachancea thermotolerans* | 2. T24 | H | 2020 |
| CR-2-12 | *Lachancea thermotolerans* | 2. T24 | H | 2020 |
| CR-2-13 | *Lachancea thermotolerans* | 2. T24 | H | 2020 |
| CR-2-15 | *Lachancea thermotolerans* | 2. T24 | H | 2020 |
| CR-2-17 | *Lachancea thermotolerans* | 2. T24 | H | 2020 |
| CR-2-18 | *Lachancea thermotolerans* | 2. T24 | H | 2020 |

| | | | | |
|---|---|---|---|---|
| CR-2-19 | *Lachancea thermotolerans* | 2. T24 | H | 2020 |
| CR-2-20 | *Lachancea thermotolerans* | 2. T24 | H | 2020 |
| CR-3-11 | *Lachancea thermotolerans* | 3. D1040 | H | 2020 |
| CR-3-12 | *Lachancea thermotolerans* | 3. D1040 | H | 2020 |
| CR-3-13 | *Lachancea thermotolerans* | 3. D1040 | H | 2020 |
| CR-3-14 | *Lachancea thermotolerans* | 3. D1040 | H | 2020 |
| CR-3-15 | *Lachancea thermotolerans* | 3. D1040 | H | 2020 |
| CR-4-13 | *Lachancea thermotolerans* | 4. D999 | H | 2020 |
| CR-4-14 | *Lachancea thermotolerans* | 4. D999 | H | 2020 |
| CR-4-15 | *Lachancea thermotolerans* | 4. D999 | H | 2020 |
| CR-501 | *Lachancea thermolerans* | 1. T0 | H | 2021 |
| CR-512 | *Lachancea thermotolerans* | 1. T0 | H | 2021 |
| CR-517 | *Lachancea thermotolerans* | 1. T0 | H | 2021 |
| CR-520 | *Lachancea thermolerans* | 1. T0 | H | 2021 |
| CR-605 | *Lachancea thermolerans* | 2. T24 | H | 2021 |
| CR-609 | *Lachancea thermolerans* | 2. T24 | H | 2021 |
| CR-618 | *Lachancea thermolerans* | 2. T24 | H | 2021 |
| CR-620 | *Lachancea thermolerans* | 2. T24 | H | 2021 |
| CR-621 | *Lachancea thermolerans* | 2. T24 | H | 2021 |
| CR-705 | *Lachancea thermolerans* | 3. D1040 | H | 2021 |
| CR-706 | *Lachancea thermolerans* | 3. D1040 | H | 2021 |
| CR-707 | *Lachancea thermolerans* | 3. D1040 | H | 2021 |
| CR-708 | *Lachancea thermolerans* | 3. D1040 | H | 2021 |
| CR-709 | *Lachancea thermolerans* | 3. D1040 | H | 2021 |
| CR-710 | *Lachancea thermolerans* | 3. D1040 | H | 2021 |
| CR-714 | *Lachancea thermolerans* | 3. D1040 | H | 2021 |
| CR-716 | *Lachancea thermolerans* | 3. D1040 | H | 2021 |
| CR-717 | *Lachancea thermolerans* | 3. D1040 | H | 2021 |
| CR-720 | *Lachancea thermolerans* | 3. D1040 | H | 2021 |
| DN-3-11 | *Lachancea thermotolerans* | 3. D1040 | H | 2020 |
| DN-3-12 | *Lachancea thermotolerans* | 3. D1040 | H | 2020 |
| DN-3-14 | *Lachancea thermotolerans* | 3. D1040 | H | 2020 |
| DN-4-14 | *Lachancea thermotolerans* | 4. D999 | H | 2020 |
| DN-4-15 | *Lachancea thermotolerans* | 4. D999 | H | 2020 |
| LS-1-03 | *Lachancea thermotolerans* | 1. T0 | H | 2020 |
| LS-1-10 | *Lachancea thermotolerans* | 1. T0 | H | 2020 |
| LS-1-15 | *Lachancea thermotolerans* | 1. T0 | H | 2020 |
| LS-1-16 | *Lachancea thermotolerans* | 1. T0 | H | 2020 |
| LS-1-17 | *Lachancea thermotolerans* | 1. T0 | H | 2020 |
| LS-2-01 | *Lachancea thermotolerans* | 2. T24 | H | 2020 |
| LS-2-04 | *Lachancea thermotolerans* | 2. T24 | H | 2020 |
| LS-2-05 | *Lachancea thermotolerans* | 2. T24 | H | 2020 |
| LS-2-06 | *Lachancea thermotolerans* | 2. T24 | H | 2020 |
| LS-2-09 | *Lachancea thermotolerans* | 2. T24 | H | 2020 |
| LS-2-12 | *Lachancea thermotolerans* | 2. T24 | H | 2020 |
| LS-2-14 | *Lachancea thermotolerans* | 2. T24 | H | 2020 |
| LS-2-15 | *Lachancea thermotolerans* | 2. T24 | H | 2020 |
| LS-2-16 | *Lachancea thermotolerans* | 2. T24 | H | 2020 |
| LS-2-18 | *Lachancea thermotolerans* | 2. T24 | H | 2020 |
| LS-2-19 | *Lachancea thermotolerans* | 2. T24 | H | 2020 |

| | | | | |
|---|---|---|---|---|
| LS-2-21 | *Lachancea thermotolerans* | 2. T24 | H | 2020 |
| LS-2-23 | *Lachancea thermotolerans* | 2. T24 | H | 2020 |
| LS-3-11 | *Lachancea thermotolerans* | 3. D1040 | H | 2020 |
| LS-3-12 | *Lachancea thermotolerans* | 3. D1040 | H | 2020 |
| LS-3-13 | *Lachancea thermotolerans* | 3. D1040 | H | 2020 |
| LS-3-14 | *Lachancea thermotolerans* | 3. D1040 | H | 2020 |
| LS-3-15 | *Lachancea thermotolerans* | 3. D1040 | H | 2020 |
| LS-4-12 | *Lachancea thermotolerans* | 4. D999 | H | 2020 |
| LS-4-13 | *Lachancea thermotolerans* | 4. D999 | H | 2020 |
| LS-4-14 | *Lachancea thermotolerans* | 4. D999 | H | 2020 |
| LS-4-15 | *Lachancea thermotolerans* | 4. D999 | H | 2020 |
| LS-507 | *Lachancea thermotolerans* | 1. T0 | H | 2021 |
| LS-508 | *Lachancea thermotolerans* | 1. T0 | H | 2021 |
| LS-517 | *Lachancea thermotolerans* | 1. T0 | H | 2021 |
| LS-520 | *Lachancea thermotolerans* | 1. T0 | H | 2021 |
| LS-522 | *Lachancea thermotolerans* | 1. T0 | H | 2021 |
| LS-618 | *Lachancea thermolerans* | 2. T24 | H | 2021 |
| LS-704 | *Lachancea thermotolerans* | 3. D1040 | H | 2021 |
| LS-706 | *Lachancea thermolerans* | 3. D1040 | H | 2021 |
| LS-716 | *Lachancea thermotolerans* | 3. D1040 | H | 2021 |
| LS-717 | *Lachancea thermotolerans* | 3. D1040 | H | 2021 |
| LS-718 | *Lachancea thermotolerans* | 3. D1040 | H | 2021 |
| MJ-1-01 | *Lachancea thermotolerans* | 1. T0 | H | 2020 |
| MJ-1-02 | *Lachancea thermotolerans* | 1. T0 | H | 2020 |
| MJ-1-04 | *Lachancea thermotolerans* | 1. T0 | H | 2020 |
| MJ-1-06 | *Lachancea thermotolerans* | 1. T0 | H | 2020 |
| MJ-1-07 | *Lachancea thermotolerans* | 1. T0 | H | 2020 |
| MJ-1-08 | *Lachancea thermotolerans* | 1. T0 | H | 2020 |
| MJ-1-11 | *Lachancea thermotolerans* | 1. T0 | H | 2020 |
| MJ-1-13 | *Lachancea thermotolerans* | 1. T0 | H | 2020 |
| MJ-1-14 | *Lachancea thermotolerans* | 1. T0 | H | 2020 |
| MJ-1-16 | *Lachancea thermotolerans* | 1. T0 | H | 2020 |
| MJ-1-17 | *Lachancea thermotolerans* | 1. T0 | H | 2020 |
| MJ-1-18 | *Lachancea thermotolerans* | 1. T0 | H | 2020 |
| MJ-1-21 | *Lachancea thermotolerans* | 1. T0 | H | 2020 |
| MJ-1-22 | *Lachancea thermotolerans* | 1. T0 | H | 2020 |
| MJ-1-23 | *Lachancea thermotolerans* | 1. T0 | H | 2020 |
| MJ-2-01 | *Lachancea thermotolerans* | 2. T24 | H | 2020 |
| MJ-2-02 | *Lachancea thermotolerans* | 2. T24 | H | 2020 |
| MJ-2-03 | *Lachancea thermotolerans* | 2. T24 | H | 2020 |
| MJ-2-04 | *Lachancea thermotolerans* | 2. T24 | H | 2020 |
| MJ-2-05 | *Lachancea thermotolerans* | 2. T24 | H | 2020 |
| MJ-2-06 | *Lachancea thermotolerans* | 2. T24 | H | 2020 |
| MJ-2-07 | *Lachancea thermotolerans* | 2. T24 | H | 2020 |
| MJ-2-10 | *Lachancea thermotolerans* | 2. T24 | H | 2020 |
| MJ-2-12 | *Lachancea thermotolerans* | 2. T24 | H | 2020 |
| MJ-2-13 | *Lachancea thermotolerans* | 2. T24 | H | 2020 |
| MJ-2-14 | *Lachancea thermotolerans* | 2. T24 | H | 2020 |
| MJ-2-15 | *Lachancea thermotolerans* | 2. T24 | H | 2020 |
| MJ-2-17 | *Lachancea thermotolerans* | 2. T24 | H | 2020 |

| | | | | |
|---|---|---|---|---|
| MJ-2-18 | *Lachancea thermotolerans* | 2. T24 | H | 2020 |
| MJ-2-20 | *Lachancea thermotolerans* | 2. T24 | H | 2020 |
| MJ-3-11 | *Lachancea thermotolerans* | 3. D1040 | H | 2020 |
| MJ-3-12 | *Lachancea thermotolerans* | 3. D1040 | H | 2020 |
| MJ-3-13 | *Lachancea thermotolerans* | 3. D1040 | H | 2020 |
| MJ-3-14 | *Lachancea thermotolerans* | 3. D1040 | H | 2020 |
| MJ-3-15 | *Lachancea thermotolerans* | 3. D1040 | H | 2020 |
| MJ-3-16 | *Lachancea thermotolerans* | 3. D1040 | H | 2020 |
| MJ-3-17 | *Lachancea thermotolerans* | 3. D1040 | H | 2020 |
| MJ-3-21 | *Lachancea thermotolerans* | 3. D1040 | H | 2020 |
| MJ-4-11 | *Lachancea thermotolerans* | 4. D999 | H | 2020 |
| MJ-4-13 | *Lachancea thermotolerans* | 4. D999 | H | 2020 |
| MJ-4-14 | *Lachancea thermotolerans* | 4. D999 | H | 2020 |
| MJ-4-15 | *Lachancea thermotolerans* | 4. D999 | H | 2020 |
| MJ-4-16 | *Lachancea thermotolerans* | 4. D999 | H | 2020 |
| MJ-4-17 | *Lachancea thermotolerans* | 4. D999 | H | 2020 |
| MJ-4-18 | *Lachancea thermotolerans* | 4. D999 | H | 2020 |
| MJ-4-19 | *Lachancea thermotolerans* | 4. D999 | H | 2020 |
| MJ-4-20 | *Lachancea thermotolerans* | 4. D999 | H | 2020 |
| MJ-4-21 | *Lachancea thermotolerans* | 4. D999 | H | 2020 |
| MJ-4-22 | *Lachancea thermotolerans* | 4. D999 | H | 2020 |
| MJ-4-23 | *Lachancea thermotolerans* | 4. D999 | H | 2020 |
| MJ-4-25 | *Lachancea thermotolerans* | 4. D999 | H | 2020 |
| MJ-4-26 | *Lachancea thermotolerans* | 4. D999 | H | 2020 |
| MJ-4-27 | *Lachancea thermotolerans* | 4. D999 | H | 2020 |
| MJ-4-28 | *Lachancea thermotolerans* | 4. D999 | H | 2020 |
| MJ-4-29 | *Lachancea thermotolerans* | 4. D999 | H | 2020 |
| MJ-4-30 | *Lachancea thermotolerans* | 4. D999 | H | 2020 |
| MJ-508 | *Lachancea thermotolerans* | 1. T0 | H | 2021 |
| MJ-518 | *Lachancea thermotolerans* | 1. T0 | H | 2021 |
| MJ-601 | *Lachancea thermotolerans* | 2. T24 | H | 2021 |
| MJ-603 | *Lachancea thermotolerans* | 2. T24 | H | 2021 |
| MJ-609 | *Lachancea thermotolerans* | 2. T24 | H | 2021 |
| MJ-618 | *Lachancea thermotolerans* | 2. T24 | H | 2021 |
| MJ-702 | *Lachancea thermotolerans* | 3. D1040 | H | 2021 |
| MJ-704 | *Lachancea thermotolerans* | 3. D1040 | H | 2021 |
| MJ-706 | *Lachancea thermotolerans* | 3. D1040 | H | 2021 |
| MJ-709 | *Lachancea thermotolerans* | 3. D1040 | H | 2021 |
| MJ-710 | *Lachancea thermotolerans* | 3. D1040 | H | 2021 |
| MJ-712 | *Lachancea thermotolerans* | 3. D1040 | H | 2021 |
| MJ-714 | *Lachancea thermotolerans* | 3. D1040 | H | 2021 |
| MJ-717 | *Lachancea thermotolerans* | 3. D1040 | H | 2021 |
| MJ-719 | *Lachancea thermotolerans* | 3. D1040 | H | 2021 |
| MJ-801 | *Lachancea thermotolerans* | 4. D999 | H | 2021 |
| MJ-811 | *Lachancea thermotolerans* | 4. D999 | H | 2021 |
| MR-4-13 | *Lachancea thermotolerans* | 4. D999 | H | 2020 |
| MR-502 | *Lachancea thermolerans* | 1. T0 | H | 2021 |
| MR-507 | *Lachancea thermolerans* | 1. T0 | H | 2021 |
| MR-519 | *Lachancea thermolerans* | 1. T0 | H | 2021 |
| MR-522 | *Lachancea thermolerans* | 1. T0 | H | 2021 |

| | | | | |
|---|---|---|---|---|
| MR-602 | *Lachancea thermolerans* | 2. T24 | H | 2021 |
| MR-605 | *Lachancea thermoolerans* | 2. T24 | H | 2021 |
| MR-611 | *Lachancea thermolerans* | 2. T24 | H | 2021 |
| MR-616 | *Lachancea thermolerans* | 2. T24 | H | 2021 |
| MR-618 | *Lachancea thermolerans* | 2. T24 | H | 2021 |
| MR-701 | *Lachancea thermolerans* | 3. D1040 | H | 2021 |
| MR-705 | *lachancea thermotolerans* | 3. D1040 | H | 2021 |
| MR-708 | *Lachancea thermolerans* | 3. D1040 | H | 2021 |
| MR-710 | *Lachancea thermotolerans* | 3. D1040 | H | 2021 |
| MR-711 | *Lachancea thermolerans* | 3. D1040 | H | 2021 |
| MR-712 | *Lachancea thermolerans* | 3. D1040 | H | 2021 |
| MR-714 | *Lachancea thermolerans* | 3. D1040 | H | 2021 |
| MR-716 | *Lachancea thermolerans* | 3. D1040 | H | 2021 |
| MR-719 | *Lachancea thermolerans* | 3. D1040 | H | 2021 |
| MR-802 | *Lachancea thermolerans* | 4. D999 | H | 2021 |
| MR-803 | *Lachancea thermolerans* | 4. D999 | H | 2021 |
| MR-804 | *Lachancea thermolerans* | 4. D999 | H | 2021 |
| MR-805 | *Lachancea thermolerans* | 4. D999 | H | 2021 |
| MR-806 | *Lachancea thermolerans* | 4. D999 | H | 2021 |
| MR-807 | *Lachancea thermolerans* | 4. D999 | H | 2021 |
| MR-809 | *Lachancea thermolerans* | 4. D999 | H | 2021 |
| MR-810 | *Lachancea thermolerans* | 4. D999 | H | 2021 |
| MR-811 | *Lachancea thermolerans* | 4. D999 | H | 2021 |
| MR-812 | *Lachancea thermolerans* | 4. D999 | H | 2021 |
| MR-813 | *Lachancea thermolerans* | 4. D999 | H | 2021 |
| MR-814 | *Lachancea thermolerans* | 4. D999 | H | 2021 |
| MR-816 | *Lachancea thermolerans* | 4. D999 | H | 2021 |
| MR-817 | *Lachancea thermolerans* | 4. D999 | H | 2021 |
| MR-819 | *Lachancea thermolerans* | 4. D999 | H | 2021 |
| MR-820 | *Lachancea thermolerans* | 4. D999 | H | 2021 |
| NG-1-03 | *Lachancea thermotolerans* | 1. T0 | P | 2020 |
| NG-1-06 | *Lachancea thermotolerans* | 1. T0 | P | 2020 |
| NG-1-08 | *Lachancea thermotolerans* | 1. T0 | P | 2020 |
| NG-1-13 | *Lachancea thermotolerans* | 1. T0 | P | 2020 |
| NG-1-16 | *Lachancea thermotolerans* | 1. T0 | P | 2020 |
| ROD21-01 | *Lachancea thermotolerans* | n.d. | ROD | 2021 |
| ROD21-02 | *Lachancea thermotolerans* | n.d. | ROD | 2021 |
| ROD21-03 | *Lachancea thermotolerans* | n.d. | ROD | 2021 |
| ROD21-04 | *Lachancea thermotolerans* | n.d. | ROD | 2021 |
| ROD21-05 | *Lachancea thermotolerans* | n.d. | ROD | 2021 |
| ROD21-06 | *Lachancea thermotolerans* | n.d. | ROD | 2021 |
| ROD21-09 | *Lachancea thermotolerans* | n.d. | ROD | 2021 |
| ROD21-10 | *Lachancea thermotolerans* | n.d. | ROD | 2021 |
| ROD21-100 | *Lachancea thermotolerans* | n.d. | ROD | 2021 |
| ROD21-101 | *Lachancea thermotolerans* | n.d. | ROD | 2021 |
| ROD21-102 | *Lachancea thermotolerans* | n.d. | ROD | 2021 |
| ROD21-103 | *Lachancea thermotolerans* | n.d. | ROD | 2021 |
| ROD21-104 | *Lachancea thermotolerans* | n.d. | ROD | 2021 |
| ROD21-105 | *Lachancea thermotolerans* | n.d. | ROD | 2021 |
| ROD21-106 | *Lachancea thermotolerans* | n.d. | ROD | 2021 |

| | | | | |
|---|---|---|---|---|
| ROD21-107 | *Lachancea thermotolerans* | n.d. | ROD | 2021 |
| ROD21-108 | *Lachancea thermotolerans* | n.d. | ROD | 2021 |
| ROD21-109 | *Lachancea thermotolerans* | n.d. | ROD | 2021 |
| ROD21-11 | *Lachancea thermotolerans* | n.d. | ROD | 2021 |
| ROD21-110 | *Lachancea thermotolerans* | n.d. | ROD | 2021 |
| ROD21-111 | *Lachancea thermotolerans* | n.d. | ROD | 2021 |
| ROD21-112 | *Lachancea thermotolerans* | n.d. | ROD | 2021 |
| ROD21-113 | *Lachancea thermotolerans* | n.d. | ROD | 2021 |
| ROD21-114 | *Lachancea thermotolerans* | n.d. | ROD | 2021 |
| ROD21-115 | *Lachancea thermotolerans* | n.d. | ROD | 2021 |
| ROD21-12 | *Lachancea thermotolerans* | n.d. | ROD | 2021 |
| ROD21-13 | *Lachancea thermotolerans* | n.d. | ROD | 2021 |
| ROD21-14 | *Lachancea thermotolerans* | n.d. | ROD | 2021 |
| ROD21-15 | *Lachancea thermotolerans* | n.d. | ROD | 2021 |
| ROD21-18 | *Lachancea thermotolerans* | n.d. | ROD | 2021 |
| ROD21-19 | *Lachancea thermotolerans* | n.d. | ROD | 2021 |
| ROD21-21 | *Lachancea thermotolerans* | n.d. | ROD | 2021 |
| ROD21-22 | *Lachancea thermotolerans* | n.d. | ROD | 2021 |
| ROD21-23 | *Lachancea thermotolerans* | n.d. | ROD | 2021 |
| ROD21-24 | *Lachancea thermotolerans* | n.d. | ROD | 2021 |
| ROD21-25 | *Lachancea thermotolerans* | n.d. | ROD | 2021 |
| ROD21-26 | *Lachancea thermotolerans* | n.d. | ROD | 2021 |
| ROD21-27 | *Lachancea thermotolerans* | n.d. | ROD | 2021 |
| ROD21-29 | *Lachancea thermotolerans* | n.d. | ROD | 2021 |
| ROD21-30 | *Lachancea thermotolerans* | n.d. | ROD | 2021 |
| ROD21-31 | *Lachancea thermotolerans* | n.d. | ROD | 2021 |
| ROD21-33 | *Lachancea thermotolerans* | n.d. | ROD | 2021 |
| ROD21-34 | *Lachancea thermotolerans* | n.d. | ROD | 2021 |
| ROD21-35 | *Lachancea thermotolerans* | n.d. | ROD | 2021 |
| ROD21-37 | *Lachancea thermotolerans* | n.d. | ROD | 2021 |
| ROD21-38 | *Lachancea thermotolerans* | n.d. | ROD | 2021 |
| ROD21-39 | *Lachancea thermotolerans* | n.d. | ROD | 2021 |
| ROD21-42 | *Lachancea thermotolerans* | n.d. | ROD | 2021 |
| ROD21-43 | *Lachancea thermotolerans* | n.d. | ROD | 2021 |
| ROD21-44 | *Lachancea thermotolerans* | n.d. | ROD | 2021 |
| ROD21-46 | *Lachancea thermotolerans* | n.d. | ROD | 2021 |
| ROD21-47 | *Lachancea thermotolerans* | n.d. | ROD | 2021 |
| ROD21-48 | *Lachancea thermotolerans* | n.d. | ROD | 2021 |
| ROD21-49 | *Lachancea thermotolerans* | n.d. | ROD | 2021 |
| ROD21-50 | *Lachancea thermotolerans* | n.d. | ROD | 2021 |
| ROD21-51 | *Lachancea thermotolerans* | n.d. | ROD | 2021 |
| ROD21-54 | *Lachancea thermotolerans* | n.d. | ROD | 2021 |
| ROD21-55 | *Lachancea thermotolerans* | n.d. | ROD | 2021 |
| ROD21-56 | *Lachancea thermotolerans* | n.d. | ROD | 2021 |
| ROD21-58 | *Lachancea thermotolerans* | n.d. | ROD | 2021 |
| ROD21-59 | *Lachancea thermotolerans* | n.d. | ROD | 2021 |
| ROD21-61 | *Lachancea thermotolerans* | n.d. | ROD | 2021 |
| ROD21-62 | *Lachancea thermotolerans* | n.d. | ROD | 2021 |
| ROD21-63 | *Lachancea thermotolerans* | n.d. | ROD | 2021 |
| ROD21-64 | *Lachancea thermotolerans* | n.d. | ROD | 2021 |

| | | | | |
|---|---|---|---|---|
| ROD21-66 | *Lachancea thermotolerans* | n.d. | ROD | 2021 |
| ROD21-67 | *Lachancea thermotolerans* | n.d. | ROD | 2021 |
| ROD21-69 | *Lachancea thermotolerans* | n.d. | ROD | 2021 |
| ROD21-70 | *Lachancea thermotolerans* | n.d. | ROD | 2021 |
| ROD21-71 | *Lachancea thermotolerans* | n.d. | ROD | 2021 |
| ROD21-72 | *Lachancea thermotolerans* | n.d. | ROD | 2021 |
| ROD21-74 | *Lachancea thermotolerans* | n.d. | ROD | 2021 |
| ROD21-75 | *Lachancea thermotolerans* | n.d. | ROD | 2021 |
| ROD21-76 | *Lachancea thermotolerans* | n.d. | ROD | 2021 |
| ROD21-77 | *Lachancea thermotolerans* | n.d. | ROD | 2021 |
| ROD21-78 | *Lachancea thermotolerans* | n.d. | ROD | 2021 |
| ROD21-80 | *Lachancea thermotolerans* | n.d. | ROD | 2021 |
| ROD21-82 | *Lachancea thermotolerans* | n.d. | ROD | 2021 |
| ROD21-83 | *Lachancea thermotolerans* | n.d. | ROD | 2021 |
| ROD21-84 | *Lachancea thermotolerans* | n.d. | ROD | 2021 |
| ROD21-86 | *Lachancea thermotolerans* | n.d. | ROD | 2021 |
| ROD21-87 | *Lachancea thermotolerans* | n.d. | ROD | 2021 |
| ROD21-88 | *Lachancea thermotolerans* | n.d. | ROD | 2021 |
| ROD21-90 | *Lachancea thermotolerans* | n.d. | ROD | 2021 |
| ROD21-91 | *Lachancea thermotolerans* | n.d. | ROD | 2021 |
| ROD21-92 | *Lachancea thermotolerans* | n.d. | ROD | 2021 |
| ROD21-93 | *Lachancea thermotolerans* | n.d. | ROD | 2021 |
| ROD21-94 | *Lachancea thermotolerans* | n.d. | ROD | 2021 |
| ROD21-96 | *Lachancea thermotolerans* | n.d. | ROD | 2021 |
| ROD21-97 | *Lachancea thermotolerans* | n.d. | ROD | 2021 |
| ROD21-99 | *Lachancea thermotolerans* | n.d. | ROD | 2021 |
| TR-1-02 | *Lachancea thermotolerans* | 1. T0 | H | 2020 |
| TR-1-08 | *Lachancea thermotolerans* | 1. T0 | H | 2020 |
| TR-1-09 | *Lachancea thermotolerans* | 1. T0 | H | 2020 |
| TR-1-16 | *Lachancea thermotolerans* | 1. T0 | H | 2020 |
| TR-2-01 | *Lachancea thermotolerans* | 2. T24 | H | 2020 |
| TR-2-12 | *Lachancea thermotolerans* | 2. T24 | H | 2020 |
| TR-3-11 | *Lachancea thermotolerans* | 3. D1040 | H | 2020 |
| TR-3-12 | *Lachancea thermotolerans* | 3. D1040 | H | 2020 |
| TR-3-13 | *Lachancea thermotolerans* | 3. D1040 | H | 2020 |
| TR-3-16 | *Lachancea thermotolerans* | 3. D1040 | H | 2020 |
| TR-3-17 | *Lachancea thermotolerans* | 3. D1040 | H | 2020 |
| TR-3-18 | *Lachancea thermotolerans* | 3. D1040 | H | 2020 |
| TR-3-19 | *Lachancea thermotolerans* | 3. D1040 | H | 2020 |
| TR-3-20 | *Lachancea thermotolerans* | 3. D1040 | H | 2020 |
| TR-4-11 | *Lachancea thermotolerans* | 4. D999 | H | 2020 |
| TR-4-12 | *Lachancea thermotolerans* | 4. D999 | H | 2020 |
| TR-4-14 | *Lachancea thermotolerans* | 4. D999 | H | 2020 |
| TR-505 | *Lachancea thermotolerans* | 1. T0 | H | 2021 |
| TR-601 | *Lachancea thermotolerans* | 2. T24 | H | 2021 |
| TR-603 | *Lachancea thermotolerans* | 2. T24 | H | 2021 |
| TR-609 | *Lachancea thermotolerans* | 2. T24 | H | 2021 |
| TR-611 | *Lachancea thermotolerans* | 2. T24 | H | 2021 |
| TR-615 | *Lachancea thermotolerans* | 2. T24 | H | 2021 |
| TR-617 | *Lachancea thermotolerans* | 2. T24 | H | 2021 |

| | | | | |
|---|---|---|---|---|
| TR-702 | *Lachancea thermotolerans* | 3. D1040 | H | 2021 |
| TR-710 | *Lachancea thermotolerans* | 3. D1040 | H | 2021 |
| TR-712 | *Lachancea thermotolerans* | 3. D1040 | H | 2021 |
| TR-718 | *Lachancea thermotolerans* | 3. D1040 | H | 2021 |
| TR-814 | *Lachancea thermotolerans* | 4. D999 | H | 2021 |
| TR-817 | *Lachancea thermotolerans* | 4. D999 | H | 2021 |
| VJ-1-02 | *Lachancea thermotolerans* | 1. T0 | P | 2020 |
| VJ-1-07 | *Lachancea thermotolerans* | 1. T0 | P | 2020 |
| VJ-1-08 | *Lachancea thermotolerans* | 1. T0 | P | 2020 |
| VJ-1-10 | *Lachancea thermotolerans* | 1. T0 | P | 2020 |
| VJ-1-13 | *Lachancea thermotolerans* | 1. T0 | P | 2020 |
| VJ-1-14 | *Lachancea thermotolerans* | 1. T0 | P | 2020 |
| VJ-1-24 | *Lachancea thermotolerans* | 1. T0 | P | 2020 |
| VJ-1-28 | *Lachancea thermotolerans* | 1. T0 | P | 2020 |
| VJ-1-34 | *Lachancea thermotolerans* | 1. T0 | P | 2020 |
| VJ-1-46 | *Lachancea thermotolerans* | 1. T0 | P | 2020 |
| VJ-1-47 | *Lachancea thermotolerans* | 1. T0 | P | 2020 |
| VJ-2-01 | *Lachancea thermotolerans* | 2. T24 | P | 2020 |
| VJ-3-11 | *Lachancea thermotolerans* | 3. D1040 | P | 2020 |
| VJ-3-15 | *Lachancea thermotolerans* | 3. D1040 | P | 2020 |
| VJ-4-11 | *Lachancea thermotolerans* | 4. D999 | P | 2020 |
| VJ-4-12 | *Lachancea thermotolerans* | 4. D999 | P | 2020 |
| VJ-4-13 | *Lachancea thermotolerans* | 4. D999 | P | 2020 |
| VJ-4-16 | *Lachancea thermotolerans* | 4. D999 | P | 2020 |
| VJ-4-17 | *Lachancea thermotolerans* | 4. D999 | P | 2020 |
| VJ-4-19 | *Lachancea thermotolerans* | 4. D999 | P | 2020 |
| VJ-4-20 | *Lachancea thermotolerans* | 4. D999 | P | 2020 |
| VJ-531 | *Lachancea thermotolerans* | 1. T0 | P | 2021 |
| VJ-542 | *Lachancea thermotolerans* | 1. T0 | P | 2021 |

**Table Supplementary 3.** Designed primers of the study for multiplex-PCR. Chromosome location, primer name, sequence and melting temperature. In bold, those employed for multilocus typing of *L. thermotolerans*

| Chromosome | Primer name | Sequence (5'→3') | Tm (ºC) |
|---|---|---|---|
| A | LTA-f/r | GTAAGAACCGCTGTAAGC | 55.0 |
| | | TACTGGATCCACCTCC | 53.4 |
| | **LTA2-f/r** | **GAGAAGAAGATGGAGTTTGGG** | **61.0** |
| | | **CTCCAGTTTCCTCGGTTCC** | **64.0** |
| B | **LTB-f/r** | **AGAAACGGGGCTTCACAGG** | **66.4** |
| | | **GTTTTGGCTAGTCCGCTTTGGG** | **64.2** |
| C | LTC-f/r | GGATTGGAGT GCGATTTGCC | 64.5 |
| | | CGCTGTAGCGATGTTTCC | 56.3 |
| D | LTD-f/r | CATCAACAGTTGTTGATGG | 58.0 |
| | | TGTGGAGGTAGATTGAGC | 56.0 |
| E | LTE-f/r | TGAGAAAACA CTGTTATGCG | 59.4 |
| | | TGTGACCAGTACGAAGGCC | 56.4 |
| F | **LTF-f/r** | **GCTCTGTCTCCACGGTGTCTGC** | **67.9** |
| | | **GGAGTTAGTGGTGGTAGAGG** | **60.5** |
| G | **LTG-f/r** | **CTAGTACTCAACCTACAACTCG** | **60.1** |
| | | **GGGACAAAGGGTAAGATTCG** | **58.40** |
| | **LTG2-f/r** | **GATAGGAAACGCTAGGAGACTCG** | **64.6** |
| | | **CTTAAGAGAACAGTCGAACTGC** | **60.1** |
| H | **LTH-f/r** | **CGAGTTTGCGGAAGACAGTGG** | **63.2** |
| | | **GATGCTAGCGCTATGACTAGC** | **61.2** |
| Mitochondrial | LTmt1-f/r | GACCCAGTTACTTATTAGGATG | 58.4 |
| | | CCATAATATTTATTATGGTAGC | 52.7 |
| | LTmt2-f/r | CATTTATAATTTATATCAAGCAG | 52.3 |
| | | CTCATTTATTAAAGGAACCC | 52.3 |
| | LTmt3-f/r | CTTCTTCTTATTTAAAAGATGC | 55.5 |
| | | CAGTTTACTGCTTTACCACTAAGC | 62.0 |
| | LTmt4-f/r | GTTTAATGGTTAAACTGTTAGATTGC | 60.1 |
| | | CTAATCATACTAAATTTAAATCACC | 55.9 |
| | LTmt5-f/r | GTATTAAAGGACAATATTCACG | 54.7 |
| | | CCTCATAAATATTTTTTATTACGG | 55.0 |

**Table Supplementary 4.** Amplicon size for each *L. thermotolerans* collection strain used in this study as positive controls. Strain code and molecular size determined using GelAnalyzer (there are as much as entries for a single isolate as number of amplicons it presents).

| strain | mw |
|---|---|
| 10-1488 | 2400 |
| 10-1488 | 1400 |
| 10-1488 | 600 |
| 10-1489 | 2500 |
| 10-1489 | 1900 |
| 10-1489 | 1400 |
| 10-1489 | 600 |
| 10-1489 | 500 |
| 10-1489 | 400 |
| 11-1806 | 2400 |
| 11-1806 | 1900 |
| 11-1806 | 1400 |
| 11-1806 | 600 |
| 11-1806 | 400 |
| 11-1806 | 300 |
| 11-1808 | 2500 |
| 11-1808 | 1800 |
| 11-1808 | 1400 |
| 11-1808 | 600 |
| 11-1808 | 400 |
| AF06 | 2200 |
| AF06 | 1300 |
| AF06 | 500 |
| AF06 | 200 |
| AF10 | 2200 |
| AF10 | 1400 |
| AF10 | 500 |
| AY13 | 2200 |
| AY13 | 1300 |
| AY13 | 1100 |
| AY13 | 500 |
| AY13 | 300 |
| CBS10519 | 2400 |
| CBS10519 | 2200 |
| CBS10519 | 2000 |
| CBS10519 | 1400 |
| CBS10519 | 600 |
| CBS10520 | 400 |
| CBS10520 | 600 |
| CBS10520 | 1400 |
| CBS10520 | 1800 |
| CBS10520 | 2500 |
| CBS10521 | 300 |
| CBS10521 | 600 |
| CBS10521 | 1200 |
| CBS10521 | 1400 |
| CBS10521 | 1800 |
| CBS10521 | 2000 |
| CBS10521 | 2200 |

| | |
|---|---|
| CBS10521 | 2400 |
| CBS137 | 600 |
| CBS137 | 800 |
| CBS137 | 1000 |
| CBS137 | 1100 |
| CBS137 | 1500 |
| CBS137 | 2200 |
| CBS1877 | 400 |
| CBS1877 | 500 |
| CBS1877 | 600 |
| CBS1877 | 900 |
| CBS1877 | 1400 |
| CBS1877 | 2000 |
| CBS1877 | 2300 |
| CBS2860 | 600 |
| CBS2860 | 1000 |
| CBS2860 | 1400 |
| CBS2860 | 2600 |
| CBS2907 | 300 |
| CBS2907 | 600 |
| CBS2907 | 1200 |
| CBS2907 | 1400 |
| CBS2907 | 2300 |
| CBS4728 | 600 |
| CBS4728 | 1400 |
| CBS4728 | 1900 |
| CBS4728 | 2300 |
| CBS5464 | 600 |
| CBS5464 | 1500 |
| CBS6467 | 300 |
| CBS6467 | 600 |
| CBS6467 | 1400 |
| CBS6467 | 2400 |
| CBS7772 | 600 |
| CBS7772 | 1400 |
| CBS7772 | 2100 |
| CBS7772 | 2300 |
| CECT 1951 | 400 |
| CECT 1951 | 600 |
| CECT 1951 | 900 |
| CECT 1951 | 1400 |
| CECT 1951 | 2200 |
| CONCERTO | 1300 |
| CONCERTO | 800 |
| CONCERTO | 500 |
| DBVPG 2551 | 300 |
| DBVPG 2551 | 600 |
| DBVPG 2551 | 1500 |
| DBVPG 2551 | 1700 |
| DBVPG 2700 | 300 |

| | |
|---|---|
| DBVPG 2700 | 500 |
| DBVPG 2700 | 1400 |
| DBVPG 2700 | 1800 |
| DBVPG 3418 | 2300 |
| DBVPG 3418 | 1500 |
| DBVPG 3418 | 1100 |
| DBVPG 3418 | 1000 |
| DBVPG 3418 | 800 |
| DBVPG 3418 | 600 |
| DBVPG 3464 | 1400 |
| DBVPG 3464 | 1000 |
| DBVPG 3464 | 500 |
| DBVPG 3466 | 300 |
| DBVPG 3466 | 600 |
| DBVPG 3466 | 1300 |
| DBVPG 3466 | 1400 |
| DBVPG 3469 | 2600 |
| DBVPG 3469 | 1400 |
| DBVPG 3469 | 600 |
| DBVPG 4014 | 2600 |
| DBVPG 4014 | 1900 |
| DBVPG 4014 | 1400 |
| DBVPG 4014 | 1300 |
| DBVPG 4014 | 600 |
| DBVPG 4035 | 2700 |
| DBVPG 4035 | 1900 |
| DBVPG 4035 | 1400 |
| DBVPG 4035 | 1300 |
| DBVPG 4035 | 600 |
| DBVPG 6867 | 2200 |
| DBVPG 6867 | 1400 |
| DBVPG 6867 | 1300 |
| DBVPG 6867 | 800 |
| DBVPG 6867 | 500 |
| DBVPG RK275 | 2100 |
| DBVPG RK275 | 600 |
| DBVPG RK361 | 2600 |
| DBVPG RK361 | 1400 |
| DBVPG RK361 | 900 |
| DBVPG RK361 | 600 |
| EM119 | 3000 |
| EM119 | 2600 |
| EM119 | 1500 |
| EM119 | 500 |
| EVEGA-128 | 1500 |
| EVEGA-128 | 1000 |
| EVEGA-128 | 600 |
| EVEGA-147 | 700 |
| EVEGA-147 | 1300 |
| EVEGA-161 | 700 |

| | |
|---|---|
| EVEGA-161 | 1300 |
| EVEGA-161 | 1500 |
| EVEGA-161 | 2100 |
| EVEGA-161.5 | 1500 |
| EVEGA-161.5 | 1300 |
| EVEGA-161.5 | 600 |
| EVEGA-161.5 | 400 |
| EVEGA-168 | 700 |
| EVEGA-168 | 1500 |
| EVEGA-168 | 1600 |
| EVEGA-171 | 400 |
| EVEGA-171 | 700 |
| EVEGA-171 | 1500 |
| EVEGA-171 | 1800 |
| EVEGA-171 | 2400 |
| EVEGA-190 | 700 |
| EVEGA-190 | 1200 |
| EVEGA-190 | 1600 |
| EVEGA-226 | 700 |
| EVEGA-226 | 1500 |
| EVEGA-231 | 600 |
| EVEGA-231 | 1500 |
| EVEGA-231 | 1800 |
| EVEGA-26 | 600 |
| EVEGA-26 | 1400 |
| EVEGA-271 | 2100 |
| EVEGA-271 | 1500 |
| EVEGA-271 | 1300 |
| EVEGA-271 | 600 |
| EVEGA-305 | 600 |
| EVEGA-305 | 1100 |
| EVEGA-305 | 1400 |
| EVEGA-305 | 2400 |
| EVEGA-307R | 600 |
| EVEGA-307R | 1100 |
| EVEGA-307R | 2100 |
| EXCELL | 300 |
| EXCELL | 500 |
| EXCELL | 1400 |
| EXCELL | 2200 |
| Fin. 89-2 | 2300 |
| Fin. 89-2 | 1900 |
| Fin. 89-2 | 1500 |
| Fin. 89-2 | 1200 |
| Fin. 89-2 | 600 |
| Fin. 89-2 | 500 |
| G13 | 2100 |
| G13 | 1400 |
| G13 | 600 |
| G32 | 2500 |

| | |
|---|---|
| G32 | 2100 |
| G32 | 1500 |
| G32 | 600 |
| ICVV43 | 2400 |
| ICVV43 | 1500 |
| ICVV43 | 1300 |
| ICVV43 | 1100 |
| ICVV43 | 500 |
| ISA2308 | 2600 |
| ISA2308 | 1400 |
| ISA2308 | 600 |
| ISA2308 | 500 |
| ISA2308 | 300 |
| ISA2380 | 2400 |
| ISA2380 | 1400 |
| ISA2380 | 1200 |
| ISA2380 | 500 |
| ISA2380 | 300 |
| KEH.34.B.3 | 1500 |
| KEH.34.B.3 | 800 |
| KEH.34.B.3 | 600 |
| LAKT | 600 |
| LAKT | 1000 |
| LAKT | 1400 |
| LAKT | 1500 |
| LEV | 600 |
| LEV | 1200 |
| LEV | 1400 |
| LG21-05 | 600 |
| LG21-05 | 1200 |
| LG21-05 | 1400 |
| MUCL31341 | 2900 |
| MUCL31341 | 2400 |
| MUCL31341 | 1400 |
| MUCL31341 | 600 |
| MUCL31341 | 500 |
| MUCL31341 | 300 |
| MY115 | 2300 |
| MY115 | 1700 |
| MY115 | 1300 |
| MY115 | 500 |
| MY115 | 300 |
| NRRL Y-27911 / IY160 | 1500 |
| NRRL Y-27911 / IY160 | 1400 |
| NRRL Y-27911 / IY160 | 800 |
| NRRL Y-27911 / IY160 | 700 |
| NRRL Y-27911 / IY160 | 500 |
| NRRL Y-27911 / IY160 | 300 |
| NRRL Y-27937 / IY38 | 1500 |
| NRRL Y-27937 / IY38 | 600 |

| | |
|---|---|
| P174 | 3100 |
| P174 | 2800 |
| P174 | 1500 |
| P174 | 600 |
| P205 | 1500 |
| P205 | 500 |
| PYCC 2908 | 2500 |
| PYCC 2908 | 1400 |
| PYCC 2908 | 500 |
| PYCC 2908 | 400 |
| PYCC 2908 | 300 |
| PYCC 4135T | 1500 |
| PYCC 4135T | 1100 |
| PYCC 4135T | 1000 |
| PYCC 4135T | 800 |
| PYCC 4135T | 600 |
| PYCC 4135T | 400 |
| PYCC 4675 | 2400 |
| PYCC 4675 | 1500 |
| PYCC 4675 | 600 |
| PYCC 6375 | 2500 |
| PYCC 6375 | 1500 |
| PYCC 6375 | 1300 |
| PYCC 6375 | 600 |
| PYCC 6806 | 1500 |
| PYCC 6806 | 500 |
| PYCC 6806 | 300 |
| PYCC 6986 | 600 |
| PYCC 6986 | 700 |
| PYCC 6986 | 1500 |
| PYCC 6986 | 1900 |
| PYCC 6986 | 2000 |
| PYCC 6986 | 2100 |
| PYCC 7194 | 300 |
| PYCC 7194 | 500 |
| PYCC 7194 | 1200 |
| PYCC 7194 | 1500 |
| PYCC 7194 | 2400 |
| PYCC 7205 | 600 |
| PYCC 7205 | 800 |
| PYCC 7205 | 1500 |
| PYCC 7205 | 1900 |
| PYCC 7205 | 2600 |
| PYCC 8789 | 600 |
| PYCC 8789 | 1100 |
| PYCC 8789 | 1400 |
| PYCC 8789 | 1500 |
| PYCC 8789 | 2200 |
| QKK | 600 |
| QKK | 1300 |

| | |
|---|---|
| QKK | 1400 |
| R21-01 | 600 |
| R21-01 | 1100 |
| R21-01 | 1400 |
| UPM 13.1 | 1400 |
| UPM 13.1 | 500 |
| UT01 | 2400 |
| UT01 | 1500 |
| UT01 | 600 |
| UT01 | 300 |
| UT09 | 1400 |
| UT09 | 500 |
| UT09 | 400 |
| UT23 | 2300 |
| UT23 | 1400 |
| UT23 | 500 |
| UT23 | 400 |
| UT23 | 300 |
| UWOPS 79-110 | 300 |
| UWOPS 79-110 | 600 |
| UWOPS 79-110 | 1500 |
| UWOPS 79-110 | 1800 |
| UWOPS 79-116 | 300 |
| UWOPS 79-116 | 400 |
| UWOPS 79-116 | 600 |
| UWOPS 79-116 | 800 |
| UWOPS 79-116 | 1500 |
| UWOPS 79-116 | 2700 |
| UWOPS 79-117 | 300 |
| UWOPS 79-117 | 600 |
| UWOPS 79-117 | 800 |
| UWOPS 79-117 | 1400 |
| UWOPS 79-117 | 1700 |
| UWOPS 79-117 | 1900 |
| UWOPS 79-162 | 500 |
| UWOPS 79-162 | 600 |
| UWOPS 79-162 | 1400 |
| UWOPS 79-162 | 1700 |
| UWOPS 79-162 | 1900 |
| UWOPS 79-164 | 600 |
| UWOPS 79-164 | 900 |
| UWOPS 79-164 | 1500 |
| UWOPS 79-164 | 1900 |
| UWOPS 79-195 | 600 |
| UWOPS 79-195 | 900 |
| UWOPS 79-195 | 1300 |
| UWOPS 79-195 | 1500 |
| UWOPS 79-248 | 400 |
| UWOPS 79-248 | 500 |
| UWOPS 79-248 | 700 |

| | |
|---|---|
| UWOPS 79-248 | 900 |
| UWOPS 79-248 | 1400 |
| UWOPS 79-248 | 1600 |
| UWOPS 79-248 | 1900 |
| UWOPS 79-248 | 2300 |
| UWOPS 79-248 | 2600 |
| UWOPS 83-1097.1 | 500 |
| UWOPS 83-1097.1 | 800 |
| UWOPS 83-1097.1 | 1400 |
| UWOPS 83-1101.1 | 500 |
| UWOPS 83-1101.1 | 800 |
| UWOPS 83-1101.1 | 1300 |
| UWOPS 83-1101.1 | 2100 |
| UWOPS 83-1101.1 | 2200 |
| UWOPS 83-1101.1 | 2400 |
| UWOPS 85-312.1 | 500 |
| UWOPS 85-312.1 | 900 |
| UWOPS 85-312.1 | 1400 |
| UWOPS 85-312.1 | 2300 |
| UWOPS 85-51.1 | 300 |
| UWOPS 85-51.1 | 500 |
| UWOPS 90-10.1 | 500 |
| UWOPS 90-10.1 | 800 |
| UWOPS 90-10.1 | 1300 |
| UWOPS 90-10.1 | 1800 |
| UWOPS 90-10.1 | 2200 |
| UWOPS 90-10.1 | 2400 |
| UWOPS 91-910.1 | 500 |
| UWOPS 91-910.1 | 1400 |
| UWOPS 91-910.1 | 1800 |
| UWOPS 91-910.1 | 2100 |
| UWOPS 91-910.1 | 2300 |
| UWOPS 91-912.1 | 500 |
| UWOPS 91-912.1 | 1500 |
| UWOPS 91-912.1 | 1800 |
| UWOPS 91-912.1 | 2100 |
| UWOPS 91-912.1 | 2400 |
| UWOPS 94-426.2 | 2500 |
| UWOPS 94-426.2 | 1800 |
| UWOPS 94-426.2 | 1400 |
| UWOPS 94-426.2 | 800 |
| UWOPS 94-426.2 | 600 |
| VINF | 600 |
| VINF | 1500 |
| VINF | 2000 |
| WN12 | 2300 |
| WN12 | 1800 |
| WN12 | 1400 |
| WN12 | 500 |
| WN13 | 2300 |

| | |
|---|---|
| WN13 | 1400 |
| WN13 | 600 |
| WN15 | 1400 |
| WN15 | 1300 |
| WN15 | 600 |
| WN15 | 1700 |
| 32SO | 2300 |
| 32SO | 2000 |
| 32SO | 1500 |
| 32SO | 600 |
| IFI 1135 | 1900 |
| IFI 1135 | 1500 |
| IFI 1135 | 1200 |
| IFI 1135 | 1000 |
| IFI 1135 | 600 |
| P46 | 2600 |
| P46 | 1800 |
| P46 | 1400 |
| P46 | 600 |
| P83 | 2100 |
| P83 | 1900 |
| P83 | 1500 |
| P83 | 600 |
| P87 | 2100 |
| P87 | 1900 |
| P87 | 1400 |
| P87 | 600 |
| P91 | 2100 |
| P91 | 1900 |
| P91 | 1400 |
| P91 | 600 |
| P91 | 300 |

**Table Supplementary 5.** Amplicon size for each *L. thermotolerans* wine related isolates used in this study. Strain code and molecular size determined using GelAnalyzer (there are as much as entries for a single isolate as number of amplicons it presents).

| strain | mw |
|---|---|
| A11-603 | 1000 |
| A11-603 | 700 |
| A11-603 | 600 |
| A11-606 | 600 |
| A11-606 | 700 |
| A11-612 | 1100 |
| A11-612 | 1000 |
| A11-612 | 600 |
| A11-617 | 600 |
| A11-617 | 700 |
| BAR20-2.02 | 700 |
| BAR20-2.02 | 1100 |
| BAR20-2.02 | 1400 |
| BAR20-2.02 | 1800 |
| BAR21-1.04 | 700 |
| BAR21-1.04 | 1400 |
| BAR21-1.04 | 1700 |
| BAR21-1.04 | 2200 |
| BAR21-1.06 | 700 |
| BAR21-1.06 | 1400 |
| BAR21-1.06 | 1700 |
| BAR21-1.06 | 2200 |
| BAR21-1.11 | 700 |
| BAR21-1.11 | 1400 |
| BAR21-1.11 | 1700 |
| BAR21-1.11 | 2200 |
| BAR21-1.12 | 700 |
| BAR21-1.12 | 1400 |
| BAR21-1.12 | 1700 |
| BAR21-1.12 | 2200 |
| BAR21-1.13 | 700 |
| BAR21-1.13 | 1400 |
| BAR21-1.13 | 1700 |
| BAR21-1.13 | 2200 |
| BAR21-1.14 | 700 |
| BAR21-1.14 | 1400 |
| BAR21-1.14 | 1700 |
| BAR21-1.14 | 2300 |
| BAR21-1.15 | 700 |
| BAR21-1.15 | 1400 |
| BAR21-1.15 | 1700 |
| BAR21-1.15 | 2300 |
| BAR21-1.18 | 700 |
| BAR21-1.18 | 1400 |
| BAR21-1.18 | 1700 |
| BAR21-1.18 | 2300 |
| BAR21-1.19 | 700 |
| BAR21-1.19 | 1400 |
| BAR21-1.19 | 1700 |

| | |
|---|---|
| BAR21-1.19 | 2300 |
| BAR21-1.20 | 700 |
| BAR21-1.20 | 1400 |
| BAR21-1.20 | 1800 |
| BAR21-1.20 | 2400 |
| BD-522 | 600 |
| BD-522 | 1400 |
| BD-522 | 1500 |
| BD-601 | 600 |
| BD-601 | 1400 |
| BD-601 | 1700 |
| BD-601 | 2100 |
| BD-612 | 600 |
| BD-612 | 1000 |
| BD-612 | 1400 |
| BD-714 | 500 |
| BD-715 | 500 |
| BD-715 | 1500 |
| BD-715 | 1600 |
| CB-109 | 600 |
| CB-109 | 1200 |
| CB-109 | 3900 |
| CB-111 | 600 |
| CB-111 | 900 |
| CB-111 | 1300 |
| CB-115 | 600 |
| CB-115 | 900 |
| CB-115 | 1300 |
| CB-115 | 1500 |
| CB-115 | 2800 |
| CB-116 | 600 |
| CB-116 | 1200 |
| CB-116 | 1500 |
| CB-116 | 2700 |
| CB-202 | 600 |
| CB-202 | 900 |
| CB-202 | 1300 |
| CB-204 | 600 |
| CB-204 | 1300 |
| CB-204 | 1600 |
| CB-205 | 500 |
| CB-205 | 1300 |
| CB-216 | 500 |
| CB-216 | 1200 |
| CB-217 | 500 |
| CB-217 | 1300 |
| CB-217 | 1500 |
| CB-217 | 2800 |
| CB-311 | 500 |
| CB-311 | 1300 |

| | |
|---|---|
| CB-311 | 1500 |
| CB-311 | 2800 |
| CB-312 | 700 |
| CB-312 | 900 |
| CB-312 | 1200 |
| CB-312 | 2800 |
| CB-313 | 700 |
| CB-313 | 1200 |
| CB-313 | 2300 |
| CB-313 | 2600 |
| CB-315 | 700 |
| CB-315 | 1200 |
| CB-316 | 700 |
| CB-317 | 600 |
| CB-317 | 1300 |
| CB-319 | 700 |
| CB-320 | 700 |
| CB-321 | 600 |
| CB-321 | 1300 |
| CB-321 | 2700 |
| CB-321 | 2800 |
| CB-506 | 600 |
| CB-507 | 500 |
| CB-507 | 1500 |
| CB-507 | 1500 |
| CB-520 | 600 |
| CB-520 | 1400 |
| CB-613 | 500 |
| CB-613 | 1400 |
| CB-613 | 1600 |
| CB-616 | 500 |
| CB-616 | 1400 |
| CB-616 | 2600 |
| CB-705 | 600 |
| CB-705 | 1400 |
| CB-705 | 1600 |
| CB-705 | 2100 |
| CB-705 | 2500 |
| CB-708 | 500 |
| CB-708 | 1300 |
| CB-708 | 1500 |
| CB-710 | 400 |
| CB-710 | 1200 |
| CB-710 | 1400 |
| CB-713 | 700 |
| CB-713 | 1100 |
| CB-713 | 1200 |
| CB-713 | 1500 |
| CB-714 | 500 |
| CB-714 | 1400 |

| | |
|---|---|
| CB-714 | 1500 |
| CR-101 | 500 |
| CR-101 | 1400 |
| CR-103 | 500 |
| CR-103 | 1400 |
| CR-104 | 500 |
| CR-104 | 1400 |
| CR-106 | 500 |
| CR-107 | 500 |
| CR-107 | 1500 |
| CR-108 | 600 |
| CR-109 | 600 |
| CR-109 | 1500 |
| CR-109 | 2100 |
| CR-109 | 2900 |
| CR-113 | 500 |
| CR-113 | 1400 |
| CR-113 | 1500 |
| CR-113 | 3000 |
| CR-115 | 700 |
| CR-116 | 600 |
| CR-116 | 1500 |
| CR-120 | 600 |
| CR-120 | 1500 |
| CR-120 | 1800 |
| CR-120 | 2000 |
| CR-120 | 2200 |
| CR-204 | 600 |
| CR-204 | 1400 |
| CR-204 | 2100 |
| CR-204 | 2300 |
| CR-210 | 600 |
| CR-211 | 600 |
| CR-212 | 600 |
| CR-213 | 600 |
| CR-215 | 600 |
| CR-215 | 1400 |
| CR-215 | 1900 |
| CR-217 | 600 |
| CR-218 | 500 |
| CR-219 | 600 |
| CR-219 | 1400 |
| CR-219 | 1700 |
| CR-219 | 2000 |
| CR-219 | 2700 |
| CR-220 | 600 |
| CR-220 | 1400 |
| CR-220 | 1900 |
| CR-311 | 500 |
| CR-311 | 1400 |

| | |
|---|---|
| CR-312 | 600 |
| CR-313 | 500 |
| CR-313 | 1400 |
| CR-314 | 500 |
| CR-315 | 600 |
| CR-315 | 1400 |
| CR-413 | 600 |
| CR-413 | 1500 |
| CR-414 | 500 |
| CR-415 | 500 |
| CR-501 | 500 |
| CR-501 | 1600 |
| CR-501 | 2200 |
| CR-520 | 500 |
| CR-520 | 1400 |
| CR-520 | 1900 |
| CR-609 | 600 |
| CR-609 | 1400 |
| CR-609 | 1500 |
| CR-609 | 1800 |
| CR-609 | 2400 |
| CR-618 | 600 |
| CR-618 | 1400 |
| CR-618 | 1700 |
| CR-618 | 1800 |
| CR-621 | 500 |
| CR-621 | 1400 |
| CR-621 | 1700 |
| CR-621 | 1900 |
| CR-621 | 2400 |
| CR-705 | 500 |
| CR-705 | 1400 |
| CR-705 | 1700 |
| CR-705 | 1800 |
| CR-705 | 2300 |
| CR-706 | 500 |
| CR-706 | 1400 |
| CR-706 | 2100 |
| CR-707 | 500 |
| CR-707 | 1400 |
| CR-707 | 1900 |
| CR-708 | 500 |
| CR-708 | 1400 |
| CR-708 | 1700 |
| CR-708 | 1800 |
| CR-708 | 2200 |
| CR-709 | 500 |
| CR-709 | 1400 |
| CR-709 | 1700 |
| CR-709 | 1900 |

| | |
|---|---|
| CR-709 | 2400 |
| CR710 | 500 |
| CR710 | 1500 |
| CR710 | 2500 |
| CR-714 | 500 |
| CR-714 | 1500 |
| CR-714 | 1600 |
| CR-714 | 2300 |
| CR-716 | 500 |
| CR-716 | 1500 |
| CR-716 | 1800 |
| CR-716 | 1900 |
| CR-717 | 600 |
| CR-717 | 1500 |
| CR-717 | 1500 |
| CR-717 | 1800 |
| CR-717 | 2900 |
| CR-720 | 500 |
| CR-720 | 1300 |
| CR-720 | 1500 |
| DN-103 | 600 |
| DN-110 | 500 |
| DN-115 | 600 |
| DN-116 | 500 |
| DN-116 | 1300 |
| DN-311 | 500 |
| DN-311 | 1300 |
| DN-311 | 1400 |
| LS-103 | 600 |
| LS-110 | 500 |
| LS-115 | 600 |
| LS-116 | 500 |
| LS-204 | 600 |
| LS-204 | 1300 |
| LS-205 | 500 |
| LS-205 | 1300 |
| LS-205 | 1600 |
| LS-205 | 2100 |
| LS-205 | 2900 |
| LS-206 | 600 |
| LS-206 | 1300 |
| LS-211 | 600 |
| LS-211 | 1300 |
| LS-212 | 500 |
| LS-214 | 500 |
| LS-214 | 1100 |
| LS-214 | 1300 |
| LS-214 | 1600 |
| LS-215 | 500 |
| LS-215 | 500 |

| | |
|---|---|
| LS-215 | 1300 |
| LS-215 | 1400 |
| LS-215 | 2700 |
| LS-216 | 500 |
| LS-216 | 1300 |
| LS-216 | 2700 |
| LS-218 | 500 |
| LS-219 | 500 |
| LS-219 | 1200 |
| LS-219 | 1200 |
| LS-219 | 2600 |
| LS-311 | 600 |
| LS-311 | 1300 |
| LS-311 | 2000 |
| LS-312 | 600 |
| LS-312 | 1400 |
| LS-312 | 2700 |
| LS-313 | 700 |
| LS-313 | 1400 |
| LS-313 | 1500 |
| LS-313 | 2700 |
| LS-315 | 600 |
| LS-315 | 1400 |
| LS-321 | 600 |
| LS-321 | 1400 |
| LS-321 | 1600 |
| LS-412 | 600 |
| LS-412 | 1400 |
| LS-412 | 1400 |
| LS-412 | 2900 |
| LS-413 | 600 |
| LS-413 | 1400 |
| LS-413 | 1400 |
| LS-413 | 2900 |
| LS-414 | 600 |
| LS-414 | 1100 |
| LS-414 | 1300 |
| LS-414 | 1500 |
| LS-415 | 600 |
| LS-415 | 1400 |
| LS-415 | 1600 |
| LS-415 | 2900 |
| LS-618 | 600 |
| LS-618 | 1400 |
| LS-618 | 2400 |
| LS-618 | 2600 |
| MJ-101 | 700 |
| MJ-101 | 1300 |
| MJ-101 | 1600 |
| MJ-104 | 700 |

| | |
|---|---|
| MJ-104 | 1600 |
| MJ-104 | 1800 |
| MJ-104 | 2300 |
| MJ-106 | 700 |
| MJ-106 | 1300 |
| MJ-106 | 1600 |
| MJ-106 | 2700 |
| MJ-107 | 600 |
| MJ-107 | 800 |
| MJ-107 | 1400 |
| MJ-107 | 1600 |
| MJ-108 | 700 |
| MJ-108 | 1600 |
| MJ-108 | 1800 |
| MJ-108 | 2100 |
| MJ-108 | 2400 |
| MJ-111 | 700 |
| MJ-111 | 1500 |
| MJ-111 | 1800 |
| MJ-116 | 600 |
| MJ-116 | 1500 |
| MJ-117 | 600 |
| MJ-117 | 1500 |
| MJ-117 | 1800 |
| MJ-117 | 2100 |
| MJ-117 | 2500 |
| MJ-118 | 500 |
| MJ-118 | 1400 |
| MJ-118 | 1900 |
| MJ-118 | 2200 |
| MJ-121 | 600 |
| MJ-121 | 1400 |
| MJ-121 | 1800 |
| MJ-121 | 2000 |
| MJ-122 | 600 |
| MJ-122 | 1400 |
| MJ-123 | 500 |
| MJ-123 | 800 |
| MJ-123 | 1400 |
| MJ-201 | 500 |
| MJ-201 | 1400 |
| MJ-202 | 700 |
| MJ-202 | 1500 |
| MJ-202 | 1700 |
| MJ-202 | 2000 |
| MJ-202 | 2300 |
| MJ-203 | 600 |
| MJ-203 | 1400 |
| MJ-203 | 1500 |
| MJ-204 | 500 |

| | |
|---|---|
| MJ-204 | 600 |
| MJ-204 | 1500 |
| MJ-204 | 2000 |
| MJ-205 | 700 |
| MJ-205 | 1400 |
| MJ-205 | 1500 |
| MJ-205 | 1600 |
| MJ-206 | 600 |
| MJ-206 | 1400 |
| MJ-206 | 1500 |
| MJ-207 | 600 |
| MJ-207 | 1500 |
| MJ-207 | 1900 |
| MJ-210 | 700 |
| MJ-210 | 1500 |
| MJ-210 | 1700 |
| MJ-210 | 2200 |
| MJ-210 | 2300 |
| MJ-212 | 600 |
| MJ-212 | 1300 |
| MJ-212 | 1500 |
| MJ-213 | 700 |
| MJ-213 | 1500 |
| MJ-213 | 1700 |
| MJ-213 | 2200 |
| MJ-213 | 2300 |
| MJ-214 | 700 |
| MJ-214 | 800 |
| MJ-214 | 1500 |
| MJ-214 | 1700 |
| MJ-214 | 2000 |
| MJ-214 | 2300 |
| MJ-215 | 700 |
| MJ-215 | 1500 |
| MJ-215 | 1900 |
| MJ-215 | 2100 |
| MJ-217 | 600 |
| MJ-217 | 1500 |
| MJ-217 | 1900 |
| MJ-217 | 2200 |
| MJ-217 | 2400 |
| MJ-218 | 600 |
| MJ-218 | 800 |
| MJ-218 | 1500 |
| MJ-220 | 600 |
| MJ-220 | 1500 |
| MJ-220 | 2100 |
| MJ-311 | 600 |
| MJ-311 | 1400 |
| MJ-311 | 1800 |

| | |
|---|---|
| MJ-311 | 2100 |
| MJ-312 | 500 |
| MJ-312 | 1400 |
| MJ-313 | 600 |
| MJ-313 | 1100 |
| MJ-313 | 1400 |
| MJ-314 | 500 |
| MJ-315 | 600 |
| MJ-315 | 1300 |
| MJ-315 | 1400 |
| MJ-315 | 2000 |
| MJ-316 | 600 |
| MJ-316 | 1400 |
| MJ-316 | 1600 |
| MJ-317 | 500 |
| MJ-317 | 1400 |
| MJ-317 | 1900 |
| MJ-317 | 2100 |
| MJ-317 | 2400 |
| MJ-321 | 600 |
| MJ-321 | 1400 |
| MJ-411 | 600 |
| MJ-413 | 500 |
| MJ-414 | 500 |
| MJ-414 | 1400 |
| MJ-414 | 2300 |
| MJ-415 | 500 |
| MJ-416 | 500 |
| MJ-417 | 500 |
| MJ-418 | 500 |
| MJ-418 | 1200 |
| MJ-418 | 1400 |
| MJ-419 | 500 |
| MJ-419 | 1400 |
| MJ-419 | 2400 |
| MJ-420 | 500 |
| MJ-420 | 1400 |
| MJ-420 | 1600 |
| MJ-421 | 500 |
| MJ-421 | 1400 |
| MJ-421 | 1900 |
| MJ-422 | 500 |
| MJ-422 | 1400 |
| MJ-422 | 2000 |
| MJ-422 | 2100 |
| MJ-422 | 2500 |
| MJ-423 | 500 |
| MJ-423 | 1400 |
| MJ-423 | 1600 |
| MJ-425 | 600 |

| | |
|---|---|
| MJ-425 | 1400 |
| MJ-425 | 1700 |
| MJ-425 | 2100 |
| MJ-426 | 600 |
| MJ-427 | 500 |
| MJ-427 | 1500 |
| MJ-428 | 500 |
| MJ-428 | 1300 |
| MJ-428 | 1500 |
| MJ-428 | 2000 |
| MJ-429 | 500 |
| MJ-429 | 1400 |
| MJ-430 | 500 |
| MJ-430 | 1400 |
| MJ-430 | 1700 |
| MJ-430 | 2100 |
| MJ-430 | 2400 |
| MJ-508 | 600 |
| MJ-508 | 1500 |
| MJ-508 | 1700 |
| MJ-508 | 2000 |
| MJ-518 | 600 |
| MJ-518 | 1500 |
| MJ-618 | 600 |
| MJ-618 | 1500 |
| MJ-618 | 1700 |
| MJ-618 | 1800 |
| MJ-618 | 2000 |
| MJ-704 | 600 |
| MJ-704 | 1100 |
| MJ-704 | 1400 |
| MJ-706 | 600 |
| MJ-706 | 1500 |
| MJ-706 | 1700 |
| MJ-706 | 2100 |
| MJ-706 | 2400 |
| MJ-709 | 500 |
| MJ-709 | 1400 |
| MJ-709 | 2000 |
| MJ-709 | 2500 |
| MJ-710 | 600 |
| MJ-710 | 1400 |
| MJ-710 | 1700 |
| MJ-710 | 2100 |
| MJ-710 | 2400 |
| MJ-801 | 500 |
| MJ-801 | 1400 |
| MJ-801 | 1700 |
| MJ-801 | 1700 |
| MJ-801 | 2400 |

| | |
|---|---|
| MJ-811 | 600 |
| MR-413 | 600 |
| MR-413 | 1300 |
| MR-502 | 600 |
| MR-502 | 1400 |
| MR-502 | 1600 |
| MR-502 | 2000 |
| MR-507 | 600 |
| MR-507 | 1500 |
| MR-507 | 1700 |
| MR-519 | 600 |
| MR-519 | 1400 |
| MR-519 | 1500 |
| MR-522 | 600 |
| MR-522 | 1500 |
| MR-522 | 1700 |
| MR-522 | 2000 |
| MR-602 | 600 |
| MR-602 | 1400 |
| MR-602 | 1500 |
| MR-611 | 600 |
| MR-611 | 1400 |
| MR-616 | 600 |
| MR-616 | 1000 |
| MR-616 | 1500 |
| MR-616 | 2000 |
| MR-616 | 2200 |
| MR-701 | 600 |
| MR-711 | 500 |
| MR-711 | 2200 |
| MR-712 | 600 |
| MR-712 | 1000 |
| MR-712 | 1400 |
| MR-714 | 400 |
| MR-719 | 600 |
| MR-719 | 1400 |
| MR-802 | 700 |
| MR-802 | 1400 |
| MR-802 | 1400 |
| MR-803 | 1400 |
| MR-803 | 1500 |
| MR-803 | 2500 |
| MR-804 | 600 |
| MR-804 | 1400 |
| MR-804 | 1900 |
| MR-804 | 2100 |
| MR-805 | 600 |
| MR-805 | 1000 |
| MR-805 | 1500 |
| MR-807 | 600 |

| | |
|---|---|
| MR-807 | 700 |
| MR-807 | 1500 |
| MR-807 | 2500 |
| MR-809 | 700 |
| MR-809 | 1400 |
| MR-809 | 1500 |
| MR-810 | 600 |
| MR-810 | 1400 |
| MR-810 | 1400 |
| MR-810 | 2600 |
| MR-811 | 600 |
| MR-811 | 1400 |
| MR-811 | 1800 |
| MR-811 | 2100 |
| MR-812 | 600 |
| MR-812 | 1400 |
| MR-812 | 1700 |
| MR-813 | 600 |
| MR-813 | 1400 |
| MR-813 | 1400 |
| MR-813 | 2600 |
| MR-816 | 500 |
| MR-816 | 1400 |
| MR-816 | 1500 |
| MR-816 | 2600 |
| MR-819 | 500 |
| MR-819 | 1500 |
| MR-819 | 1600 |
| MR-819 | 2200 |
| NG-103 | 500 |
| NG-103 | 1300 |
| NG-103 | 1700 |
| NG-106 | 400 |
| NG-106 | 1300 |
| NG-106 | 1800 |
| NG-106 | 2300 |
| NG-106 | 2600 |
| NG-108 | 400 |
| NG-108 | 1300 |
| NG-113 | 500 |
| NG-113 | 1200 |
| NG-113 | 1600 |
| NG-113 | 2000 |
| NG-116 | 500 |
| NG-116 | 1600 |
| NG-116 | 2400 |
| NG-116 | 2800 |
| R21-33 | 2200 |
| R21-33 | 2000 |
| R21-33 | 1700 |

| | |
|---|---|
| R21-33 | 1400 |
| R21-33 | 600 |
| R21-35 | 2800 |
| R21-35 | 2100 |
| R21-35 | 1600 |
| R21-35 | 600 |
| R21-35 | 100 |
| R21-97 | 2200 |
| R21-97 | 1400 |
| R21-97 | 1200 |
| R21-97 | 600 |
| ROD21-01 | 500 |
| ROD21-01 | 1000 |
| ROD21-02 | 600 |
| ROD21-02 | 1400 |
| ROD21-02 | 1600 |
| ROD21-02 | 2000 |
| ROD21-03 | 600 |
| ROD21-03 | 1400 |
| ROD21-04 | 600 |
| ROD21-04 | 1400 |
| ROD21-04 | 2000 |
| ROD21-05 | 600 |
| ROD21-05 | 1400 |
| ROD21-100 | 600 |
| ROD21-100 | 1300 |
| ROD21-100 | 1500 |
| ROD21-101 | 700 |
| ROD21-101 | 1300 |
| ROD21-101 | 1500 |
| ROD21-102 | 700 |
| ROD21-102 | 1300 |
| ROD21-102 | 1500 |
| ROD21-103 | 700 |
| ROD21-103 | 1300 |
| ROD21-103 | 1500 |
| ROD21-104 | 700 |
| ROD21-104 | 1300 |
| ROD21-104 | 1500 |
| ROD21-106 | 700 |
| ROD21-106 | 1200 |
| ROD21-106 | 1500 |
| ROD21-107 | 700 |
| ROD21-107 | 1200 |
| ROD21-107 | 1400 |
| ROD21-108 | 700 |
| ROD21-108 | 1200 |
| ROD21-108 | 1400 |
| ROD21-109 | 600 |
| ROD21-109 | 1200 |

| | |
|---|---|
| ROD21-109 | 1400 |
| ROD21-110 | 600 |
| ROD21-110 | 1200 |
| ROD21-110 | 1400 |
| ROD21-111 | 600 |
| ROD21-111 | 1200 |
| ROD21-111 | 1400 |
| ROD21-112 | 600 |
| ROD21-112 | 1200 |
| ROD21-112 | 1400 |
| ROD21-113 | 600 |
| ROD21-114 | 600 |
| ROD21-114 | 1200 |
| ROD21-114 | 1400 |
| ROD21-115 | 600 |
| ROD21-115 | 1200 |
| ROD21-115 | 1400 |
| ROD21-13 | 600 |
| ROD21-13 | 1400 |
| ROD21-14 | 500 |
| ROD21-14 | 1000 |
| ROD21-14 | 1400 |
| ROD21-15 | 600 |
| ROD21-15 | 1500 |
| ROD21-15 | 1700 |
| ROD21-15 | 1800 |
| ROD21-18 | 600 |
| ROD21-18 | 1400 |
| ROD21-18 | 1500 |
| ROD21-19 | 600 |
| ROD21-19 | 1400 |
| ROD21-20 | 600 |
| ROD21-20 | 1400 |
| ROD21-20 | 1700 |
| ROD21-22 | 600 |
| ROD21-22 | 800 |
| ROD21-22 | 1200 |
| ROD21-22 | 1400 |
| ROD21-22 | 1800 |
| ROD21-23 | 600 |
| ROD21-23 | 1400 |
| ROD21-23 | 1900 |
| ROD21-24 | 600 |
| ROD21-24 | 1500 |
| ROD21-24 | 2100 |
| ROD21-25 | 600 |
| ROD21-25 | 1500 |
| ROD21-25 | 2000 |
| ROD21-26 | 600 |
| ROD21-26 | 1500 |

| | |
|---|---|
| ROD21-26 | 1900 |
| ROD21-27 | 600 |
| ROD21-27 | 1500 |
| ROD21-27 | 1900 |
| ROD21-29 | 600 |
| ROD21-29 | 1500 |
| ROD21-30 | 600 |
| ROD21-30 | 1500 |
| ROD21-30 | 1900 |
| ROD21-31 | 500 |
| ROD21-31 | 1500 |
| ROD21-31 | 1600 |
| ROD21-33 | 600 |
| ROD21-33 | 1500 |
| ROD21-33 | 1700 |
| ROD21-33 | 2100 |
| ROD21-33 | 2300 |
| ROD21-34 | 600 |
| ROD21-34 | 1500 |
| ROD21-34 | 2000 |
| ROD21-35 | 600 |
| ROD21-35 | 1500 |
| ROD21-35 | 2000 |
| ROD21-37 | 600 |
| ROD21-37 | 1500 |
| ROD21-37 | 2300 |
| ROD21-38 | 600 |
| ROD21-38 | 1500 |
| ROD21-42 | 600 |
| ROD21-42 | 1600 |
| ROD21-46 | 600 |
| ROD21-46 | 1500 |
| ROD21-46 | 1900 |
| ROD21-46 | 2000 |
| ROD21-46 | 2500 |
| ROD21-47 | 600 |
| ROD21-47 | 1500 |
| ROD21-47 | 2000 |
| ROD21-48 | 700 |
| ROD21-48 | 1500 |
| ROD21-48 | 2100 |
| ROD21-48 | 2400 |
| ROD21-49 | 700 |
| ROD21-49 | 900 |
| ROD21-49 | 1100 |
| ROD21-49 | 1500 |
| ROD21-49 | 2100 |
| ROD21-50 | 500 |
| ROD21-50 | 700 |
| ROD21-50 | 1400 |

| | |
|---|---|
| ROD21-50 | 1700 |
| ROD21-50 | 2100 |
| ROD21-51 | 500 |
| ROD21-51 | 1400 |
| ROD21-51 | 1600 |
| ROD21-51 | 2300 |
| ROD21-56 | 600 |
| ROD21-56 | 700 |
| ROD21-56 | 1400 |
| ROD21-56 | 2100 |
| ROD21-56 | 2500 |
| ROD21-58 | 500 |
| ROD21-58 | 1400 |
| ROD21-61 | 600 |
| ROD21-61 | 1400 |
| ROD21-61 | 1700 |
| ROD21-61 | 2500 |
| ROD21-62 | 700 |
| ROD21-62 | 700 |
| ROD21-62 | 1400 |
| ROD21-63 | 500 |
| ROD21-63 | 1400 |
| ROD21-63 | 1900 |
| ROD21-63 | 2000 |
| ROD21-64 | 700 |
| ROD21-64 | 1400 |
| ROD21-64 | 2100 |
| ROD21-66 | 700 |
| ROD21-66 | 1400 |
| ROD21-66 | 2100 |
| ROD21-67 | 700 |
| ROD21-67 | 1400 |
| ROD21-67 | 1700 |
| ROD21-69 | 700 |
| ROD21-69 | 700 |
| ROD21-69 | 1400 |
| ROD21-69 | 2200 |
| ROD21-70 | 700 |
| ROD21-70 | 1400 |
| ROD21-70 | 2100 |
| ROD21-71 | 700 |
| ROD21-71 | 700 |
| ROD21-71 | 1400 |
| ROD21-72 | 600 |
| ROD21-72 | 1400 |
| ROD21-74 | 600 |
| ROD21-74 | 1400 |
| ROD21-75 | 600 |
| ROD21-75 | 700 |
| ROD21-75 | 1300 |

| | |
|---|---|
| ROD21-76 | 600 |
| ROD21-76 | 700 |
| ROD21-76 | 1300 |
| ROD21-77 | 600 |
| ROD21-77 | 700 |
| ROD21-77 | 1300 |
| ROD21-78 | 600 |
| ROD21-78 | 700 |
| ROD21-78 | 1300 |
| ROD21-80 | 500 |
| ROD21-80 | 1300 |
| ROD21-83 | 600 |
| ROD21-83 | 700 |
| ROD21-83 | 1300 |
| ROD21-87 | 600 |
| ROD21-87 | 1600 |
| ROD21-87 | 1800 |
| ROD21-90 | 500 |
| ROD21-90 | 1600 |
| ROD21-90 | 2200 |
| ROD21-91 | 600 |
| ROD21-91 | 1600 |
| ROD21-91 | 2100 |
| ROD21-92 | 500 |
| ROD21-92 | 1600 |
| ROD21-93 | 600 |
| ROD21-93 | 1600 |
| ROD21-93 | 2200 |
| ROD21-94 | 500 |
| ROD21-94 | 1600 |
| ROD21-94 | 2200 |
| ROD21-97 | 600 |
| ROD21-97 | 1200 |
| ROD21-97 | 1400 |
| ROD21-99 | 600 |
| TR-102 | 500 |
| TR-102 | 1400 |
| TR-102 | 1600 |
| TR-108 | 600 |
| TR-108 | 1400 |
| TR-108 | 1700 |
| TR-108 | 1900 |
| TR-109 | 600 |
| TR-109 | 1300 |
| TR-109 | 1400 |
| TR-116 | 500 |
| TR-116 | 700 |
| TR-116 | 1400 |
| TR-116 | 1800 |
| TR-116 | 2200 |

| | |
|---|---|
| TR-116 | 2400 |
| TR-201 | 600 |
| TR-201 | 1300 |
| TR-201 | 1400 |
| TR-201 | 2600 |
| TR-212 | 600 |
| TR-212 | 1400 |
| TR-212 | 1700 |
| TR-212 | 1900 |
| TR-212 | 2700 |
| TR-311 | 500 |
| TR-311 | 800 |
| TR-312 | 500 |
| TR-312 | 1300 |
| TR-312 | 1400 |
| TR-313 | 600 |
| TR-313 | 1400 |
| TR-313 | 1800 |
| TR-313 | 2000 |
| TR-313 | 2800 |
| TR-316 | 600 |
| TR-316 | 1400 |
| TR-316 | 2100 |
| TR-317 | 500 |
| TR-318 | 600 |
| TR-319 | 600 |
| TR-319 | 1500 |
| TR-319 | 2200 |
| TR-319 | 2500 |
| TR-320 | 500 |
| TR-320 | 1500 |
| TR-320 | 1700 |
| TR-320 | 2500 |
| TR-411 | 500 |
| TR-411 | 1500 |
| TR-412 | 400 |
| TR-412 | 800 |
| TR-412 | 1500 |
| TR-412 | 2600 |
| TR-414 | 500 |
| TR-414 | 1500 |
| TR-505 | 500 |
| TR-505 | 1500 |
| TR-505 | 1600 |
| TR-505 | 1900 |
| TR-505 | 2500 |
| TR-601 | 600 |
| TR-601 | 1400 |
| TR-601 | 2400 |
| TR-609 | 600 |

| | |
|---|---|
| TR-611 | 500 |
| TR-615 | 500 |
| TR-615 | 1400 |
| TR-615 | 1700 |
| TR-617 | 600 |
| TR-702 | 600 |
| TR-710 | 600 |
| TR-710 | 1400 |
| TR-710 | 1600 |
| TR-710 | 2200 |
| TR-712 | 600 |
| TR-712 | 1500 |
| TR-712 | 1800 |
| TR-712 | 1900 |
| TR-712 | 2100 |
| TR-718 | 600 |
| TR-718 | 1500 |
| TR-718 | 1700 |
| TR-718 | 2000 |
| TR-718 | 2400 |
| TR-814 | 600 |
| VJ-102 | 600 |
| VJ-102 | 800 |
| VJ-102 | 1200 |
| VJ-102 | 1300 |
| VJ-102 | 1700 |
| VJ-102 | 2500 |
| VJ-107 | 600 |
| VJ-107 | 1300 |
| VJ-107 | 1900 |
| VJ-108 | 600 |
| VJ-108 | 1000 |
| VJ-108 | 1400 |
| VJ-108 | 1800 |
| VJ-110 | 600 |
| VJ-110 | 1000 |
| VJ-110 | 1400 |
| VJ-110 | 1900 |
| VJ-110 | 2200 |
| VJ-113 | 600 |
| VJ-113 | 900 |
| VJ-114 | 600 |
| VJ-114 | 1100 |
| VJ-114 | 1400 |
| VJ-114 | 1700 |
| VJ-124 | 600 |
| VJ-124 | 1200 |
| VJ-124 | 1400 |
| VJ-124 | 2100 |
| VJ-128 | 600 |

| | |
|---|---|
| VJ-128 | 1000 |
| VJ-128 | 1400 |
| VJ-128 | 1500 |
| VJ-134 | 600 |
| VJ-134 | 1400 |
| VJ-134 | 2100 |
| VJ-146 | 600 |
| VJ-146 | 900 |
| VJ-146 | 1300 |
| VJ-147 | 600 |
| VJ-147 | 1400 |
| VJ-208 | 600 |
| VJ-208 | 1500 |
| VJ-311 | 600 |
| VJ-311 | 1500 |
| VJ-311 | 2200 |
| VJ-411 | 600 |
| VJ-411 | 1100 |
| VJ-411 | 1600 |
| VJ-411 | 1700 |
| VJ-412 | 600 |
| VJ-412 | 1100 |
| VJ-412 | 1600 |
| VJ-412 | 1700 |
| VJ-413 | 600 |
| VJ-413 | 1100 |
| VJ-413 | 1500 |
| VJ-413 | 1700 |
| VJ-416 | 600 |
| VJ-416 | 1600 |
| VJ-416 | 2900 |
| VJ-417 | 600 |
| VJ-417 | 1100 |
| VJ-417 | 1500 |
| VJ-417 | 1700 |
| VJ-419 | 600 |
| VJ-419 | 1100 |
| VJ-419 | 1500 |
| VJ-419 | 1700 |
| VJ-420 | 600 |
| VJ-420 | 1000 |
| VJ-420 | 1500 |
| VJ-517 | 600 |
| VJ-517 | 1100 |
| VJ-517 | 1400 |
| VJ-517 | 1900 |
| VJ-525 | 600 |
| VJ-525 | 1400 |
| VJ-525 | 2500 |
| VJ-624 | 300 |

**Figure Supplementary 1.** Correlation analysis between geographical and genotypic distance. **A)** Collection strains. **B)** Isolated strains.

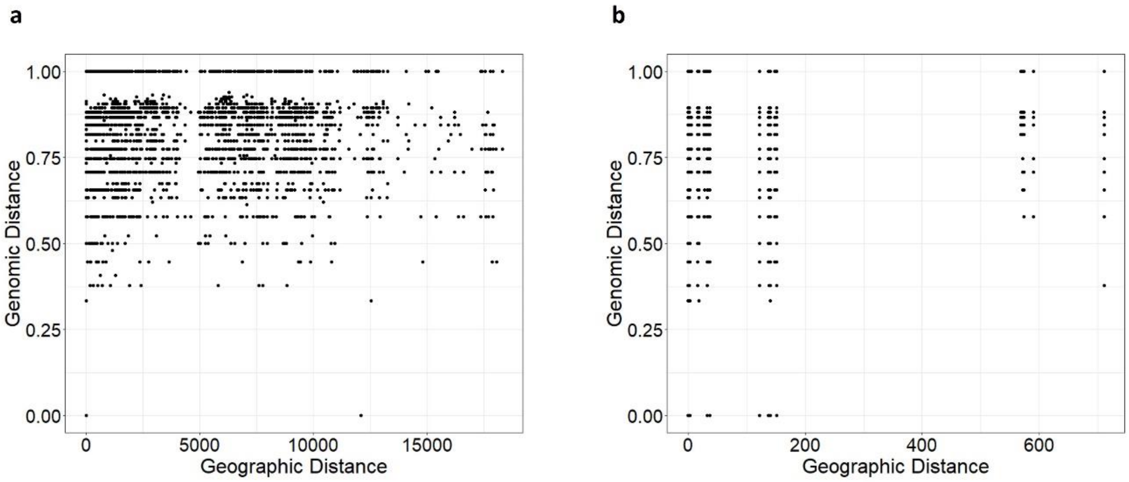

**Figure Supplementary 2.** Dendrogram showing the different clusters based on Sørensen-Dice coefficient constructed using Ward's methods for *L. thermotolerans* natural isolates.

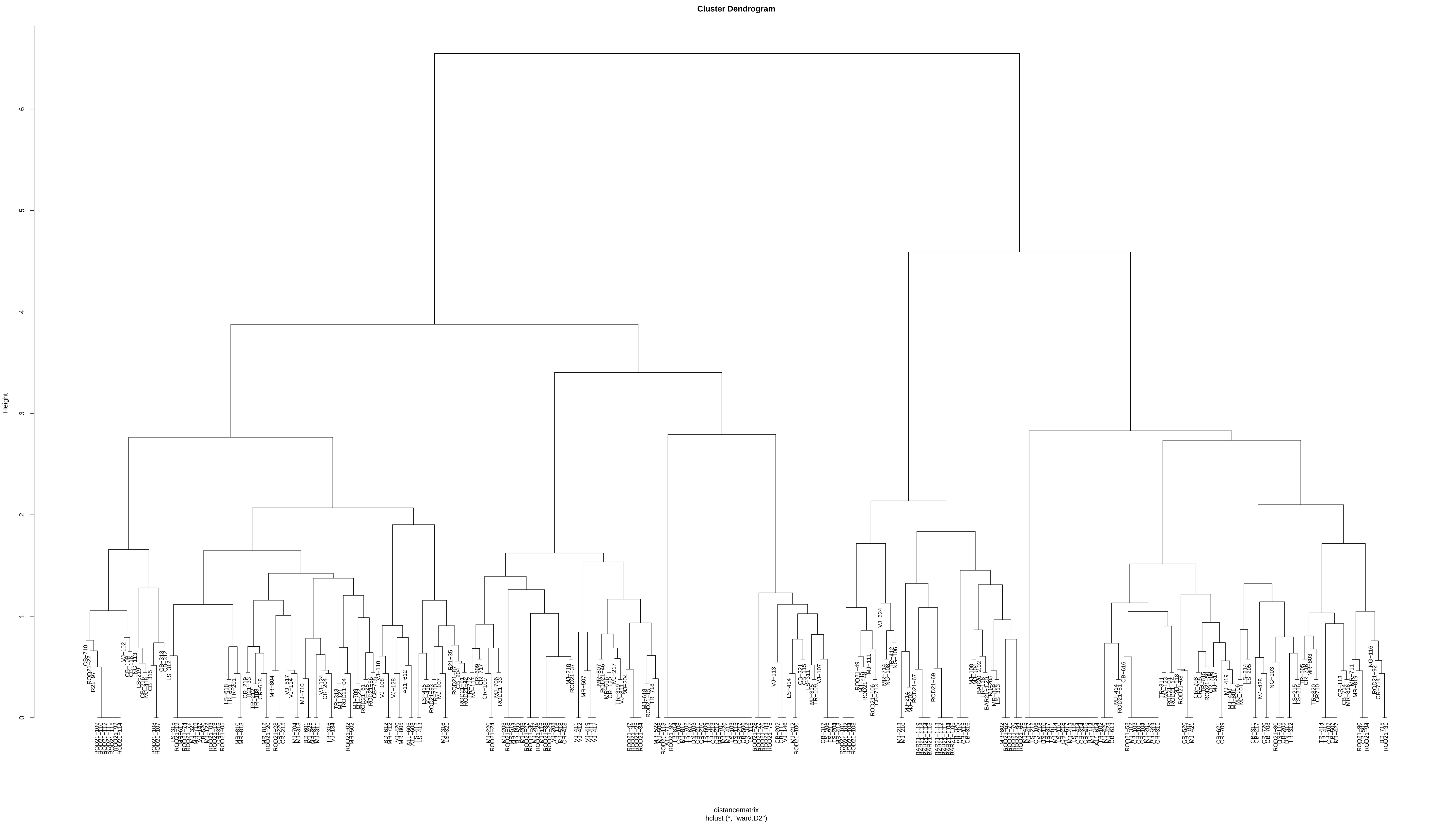

Cluster Dendrogram
Height
0
1
2
3
4
5
6
distancematrix
hclust (*, "ward.D2")
